# Importance of Glutamine in Synaptic Vesicles Revealed by Functional Studies of SLC6A17 and Its Mutations Pathogenic for Intellectual Disability

**DOI:** 10.1101/2022.10.25.513688

**Authors:** Xiaobo Jia, Jiemin Zhu, Xiling Bian, Sulin Liu, Sihan Yu, Wenjun Liang, Lifen Jiang, Renbo Mao, Yi Rao

## Abstract

Human mutations in the gene encoding the solute carrier (SLC) 6A17 caused intellectual disability (ID). The physiological role of SLC6A17 and pathogenesis of Slc6a17-based-ID were both unclear. Here we report learning deficits in SLC6A17 knockout and point mutants. Biochemistry, proteomics and electron microscopy (EM) support SLC6A17 protein localization in synaptic vesicles (SVs). Chemical analysis of SVs by liquid chromatography coupled to mass spectrometry (LC-MS) revealed glutamine (Gln) in SVs containing SLC6A17. Virally mediated overexpression of SLC6A17 increased Gln in SVs. Either genetic or virally mediated targeting of SLC6A17 reduced Gln in SVs. One ID mutation caused SLC6A17 mislocalization while the other caused defective Gln transport. Multidisciplinary approaches with 7 types of genetically modified mice have shown Gln as an endogenous substrate of SLC 6A17, uncovered Gln as a new molecule in SVs, established the necessary and sufficient roles of SLC6A17 in Gln transport into SVs, and suggested SV Gln decrease as the key pathogenetic mechanism in human ID.

## Introduction

With an approximate prevalence of 1%, intellectual disability (ID) in humans is a neurodevelopmental disorder with debilitating effects on patients (Maulik et al., 2011). Molecular genetic research into ID has identified multiple genes whose defects underly ID (de Ligt et al., 2012; Gilissen et al., 2014; Hamdan et al., 2014; Hu et al., 2019; Khan et al., 2016; Lelieveld et al., 2016; Rauch et al., 2012; Ropers, 2010; Vissers et al., 2016). Functional studies of products encoded by these genes are essential for our mechanistic understanding of ID pathogenesis.

Mutations causing ID were found in the gene encoding SLC6A17 (Iqbal et al., 2015; Waltl, 2015). SLC6A17, also known as Rxt1, NTT4, XT1, or B^0^AT3 (el Mestikawy et al., 1994; Liu et al., 1993), was discovered 29 years ago (el Mestikawy *et al*., 1994; Liu *et al*., 1993). It is predominantly expressed in the nervous system (el Mestikawy *et al*., 1994; Fischer et al., 1999; Hagglund et al., 2013; Jursky and Nelson, 1999; Kachidian, 1999; Luque et al., 1996; Masson et al., 1995; Masson et al., 1999). SLC6A17 protein was localized on SVs (Fischer *et al*., 1999; Kachidian, 1999; Masson *et al*., 1999). Those facts, together with its membership in the SLC6 or the neurotransmitter transporter (NTT) family, suggest that SLC6A17 could be a vesicular transporter for a neurotransmitter(s).

Early efforts failed to identify substrates transported by SLC6A17 (Broer, 2006; el Mestikawy *et al*., 1994; Liu *et al*., 1993). A later study using pheochromocytoma (PC)12 cells and SLC6A17-tranfected Chinese hamster ovary (CHO) cells found that SLC6A17 could transport 4 amino acids (AA): proline (Pro), glycine (Gly), leucine (Leu) and alanine (Ala) (Parra et al., 2008). Another study using human embryonic kidney (HEK) cells reported transport of 9 AAs: Leu, methionine (Met), Pro, cysteine (Cys), Ala, glutamine (Gln), serine (Ser), histidine (His) and Gly (Zaia and Reimer, 2009). There are three differences between these two studies: that the second study reported 5 more amino acids (Met, Cys, Gln, Ser and His) as substrates of SLC6A17, that the second study used modified SLC6A17 to facilitate its membrane localization (Zaia and Reimer, 2009), and that one reported Na^+^ dependence (Zaia and Reimer, 2009) whereas the other reported H^+^ dependence (Parra *et al*., 2008). However, none of these AAs have been found in the SVs. It remains unknown which of the 4 or 9 AAs, if any, are present in the SVs in an SLC6A17 dependent manner *in vivo*.

Thus, how mutations in a single AA of SLC6A17 lead to ID is unknown, the substrate(s) physiologically transported by SLC6A17 is unknown and behavioral phenotypes of animals lacking SLC6A17 or carrying any mutation pathogenic in human patients have not been characterized.

We have now generated mouse mutants either lacking the Slc6a17 gene or mimicking a point mutation found in ID patients. They had similar phenotypes including deficient learning and memory, indicating that the human mutation was a loss of function (LOF) mutation. These are the first animal models of ID caused by Slc6a17 mutations. While two in vitro studies with cell lines suggested that SLC6A17 could transport up to 9 neutral AAs, we could only find that Gln was present in the SVs. From multiple gain of function (GOF) and LOF experiments, we have obtained evidence that SLC6A17 is not only sufficient for Gln presence in the SVs *in vivo*, but it is also physiologically necessary for Gln presence in the SVs. Thus, we have found the endogenous substrate for SLC6A17. Of the two known human mutations pathogenic for ID (Iqbal *et al*., 2015), we provide evidence that one caused the SLC6A17 to be mislocalized subcellularly and the other, while still on SVs, was defective in Gln transport. Decreases of Gln in SVs caused by SLC6A17 mutations were not correlated with any decrease in glutamate (Glu) or gamma-aminobutyric acid (GABA), dissociating vesicular Gln from the Glu/GABA-Gln cycle between neurons and glia. In addition to dissecting the molecular mechanisms underlying ID caused by Slc6a17 mutations, we report for the first time that Gln is robustly and reproducibly detected in the SVs, which should stimulate further research into its potential roles in neurotransmission.

## Results

### Pattern of Slc6a17 expression

To examine the pattern of Slc6a17 expression, we designed a knock-in mouse line Slc6a17^-2A-CreERT2^ (Figure 1A). These mice were generated by in frame fusion of a T2A sequence (Ahier and Jarriault, 2014; Daniels et al., 2014; Trichas et al., 2008) and CreERT2 (Feil et al., 1996; Gu et al., 1994; Indra et al., 1999; Sauer and Henderson, 1988) to the C terminus of Slc6a17 with its stop codon removed (Figure S1A). We crossed the Slc6a17^-2A-CreERT2^ mice with the Ai14 reporter line which contained floxed stop-tdTomato (Madisen et al., 2010). Treatment with Tamoxifen removed the stop codon and thus allowed specific expression of tdTomato in Slc6a17 positive cells (Figure 1A, Figure S1A). Labeled cells were exclusively neuronal with no glial expression detected.

**Figure 1.**
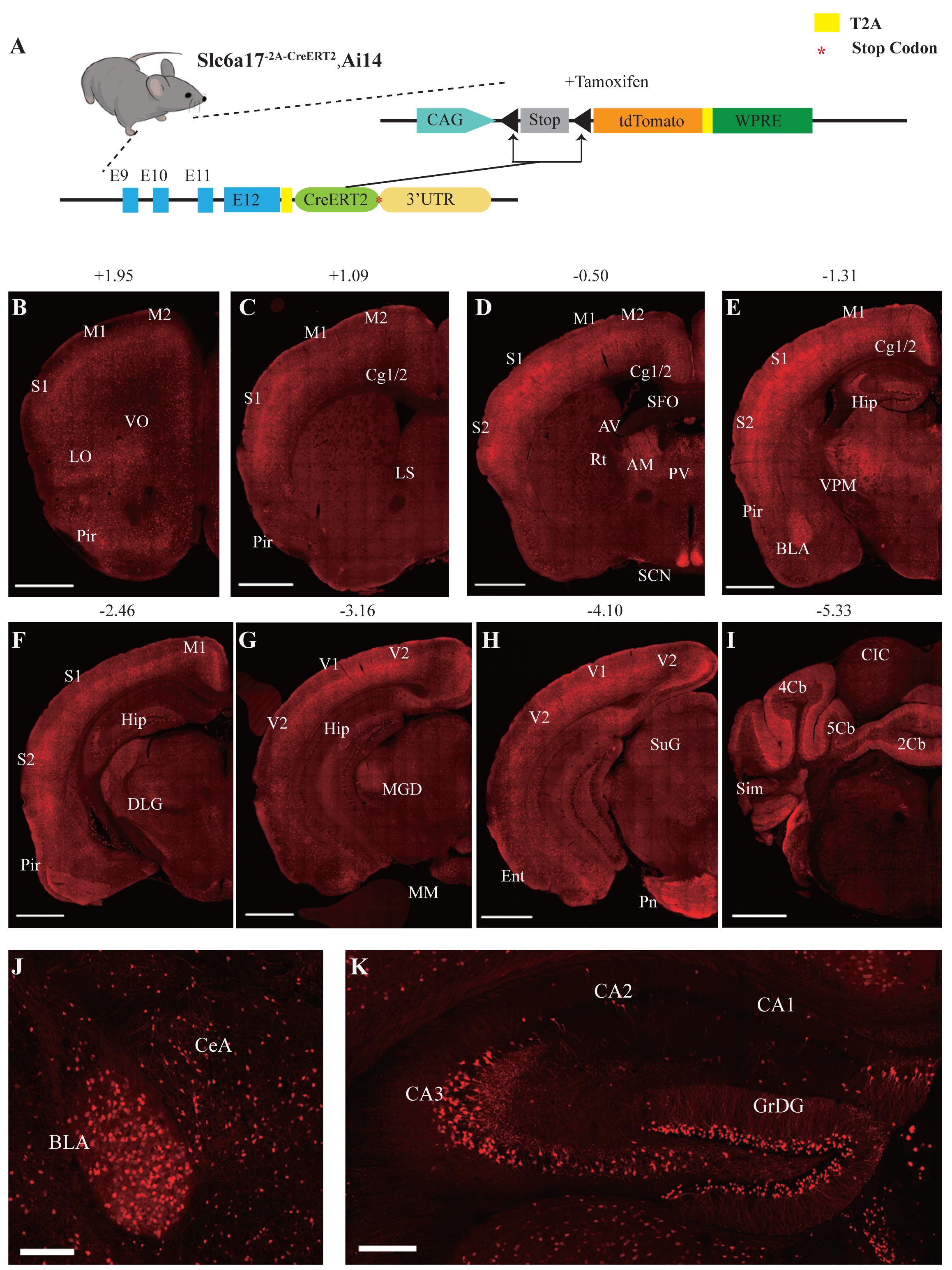
Slc6a17 Expression in the Mouse Brain. (A) A schematic diagram illustrating the strategy for the generation of Slc6a17^-2A-CreERT2^ mice. Crossing of Slc6a17^-2A-CreERT2^ mice with Ai14 (LSL-tdTomato) mice allowed specific labeling of Slc6a17-expressing neuron after tamoxifen injection. More details in Figure S1A. (B-I) Representative coronal sections of Slc6a17^-2A-CreERT2^::Ai14 mice. Numbers above images indicate the anteroposterior position of the section from Bregma in millimeters (mm), based on(Paxinos and Franklin, 2019). Scale bars=1 mm. (J) Slc6a17-positive neurons in the basolateral amygdaloid nucleus (BLA) and the central amygdaloid nucleus (CeA). Scale bars=100 μ. (K) Hippocampal expression of Slc6a17. Slc6a17-positive neurons are densely distributed in the granule cell layer of the dentate gyrus (GrDG), and CA3, with little expression in the CA1 and the CA2. Scale bars=200 μm. Abbreviations: 2Cb, lobule 2 of the cerebellar vermis; 3Cb, lobule 3 of the cerebellar vermis; 4/5Cb, lobule 4 and 5 of the cerebellar vermis; AON, accessory olfactory nucleus; AM, anteromedial thalamic nucleus; AV, anteroventral thalamic nucleus; BLA, basolateral amygdaloid nucleus; Ent, entorhinal cortex; CB, cerebellum; Cg1/2, cingulate ccortex; CIC, central nucleus of the inferior colliculus; Ctx, cortex; Hip, hippocampus; LO, lateral orbital cortex; LS, lateral septal nucleus; M1, primary motor cortex; M2, secondary motor cortex; OB, main olfactory bulb; Pir, piriform cortex; Pn, pontine nuclei; Rt, reticular thalamic nucleus; S1, primary somatosensory cortex; S2, secondary somatosensory cortex; SCN, suprachiasmatic nucleus; SFO, subfornical organ; SuG, superficial gray layer of superior colliculus; Sim, simple lobule; Th, thalamus; V1, primary visual cortex; V2, secondary visual cortex; VPM, ventral posteromedial nucleus.

Slc6a17 expression was found in the neocortex, the thalamus, the amygdala, the hippocampus, the pontine nuclei and the brainstem (Figure 1B to 1K, Figure S1B and S1C). Strong expression of Slc6a17 was detected in the dentate gyrus (DG) and the CA3 region of the hippocampus (Figure 1K), which is essential for spatial learning and memory. Little expression was observed in either CA1 or CA2 (Figure 1K). Slc6a17 was also expressed in the basolateral amygdala (BLA) (Figure 1J), a brain area essential for emotion and fear learning.

### Behavioral deficits of Slc6a17 mutant mice

To investigate the functional role of Slc6a17, we generated a knock-out (KO) mouse mutant line with exon 2 deleted from the Slc6a17 gene (Figure 2A, Figure S2A). These Slc6a17-KO mice were not significantly different from the wild type (WT, Slc6a17^+/+^) mice in body weight (Figure S3A), basal activities (Figure S3B), or short-term memory (Figure S3C).

**Figure 2.**
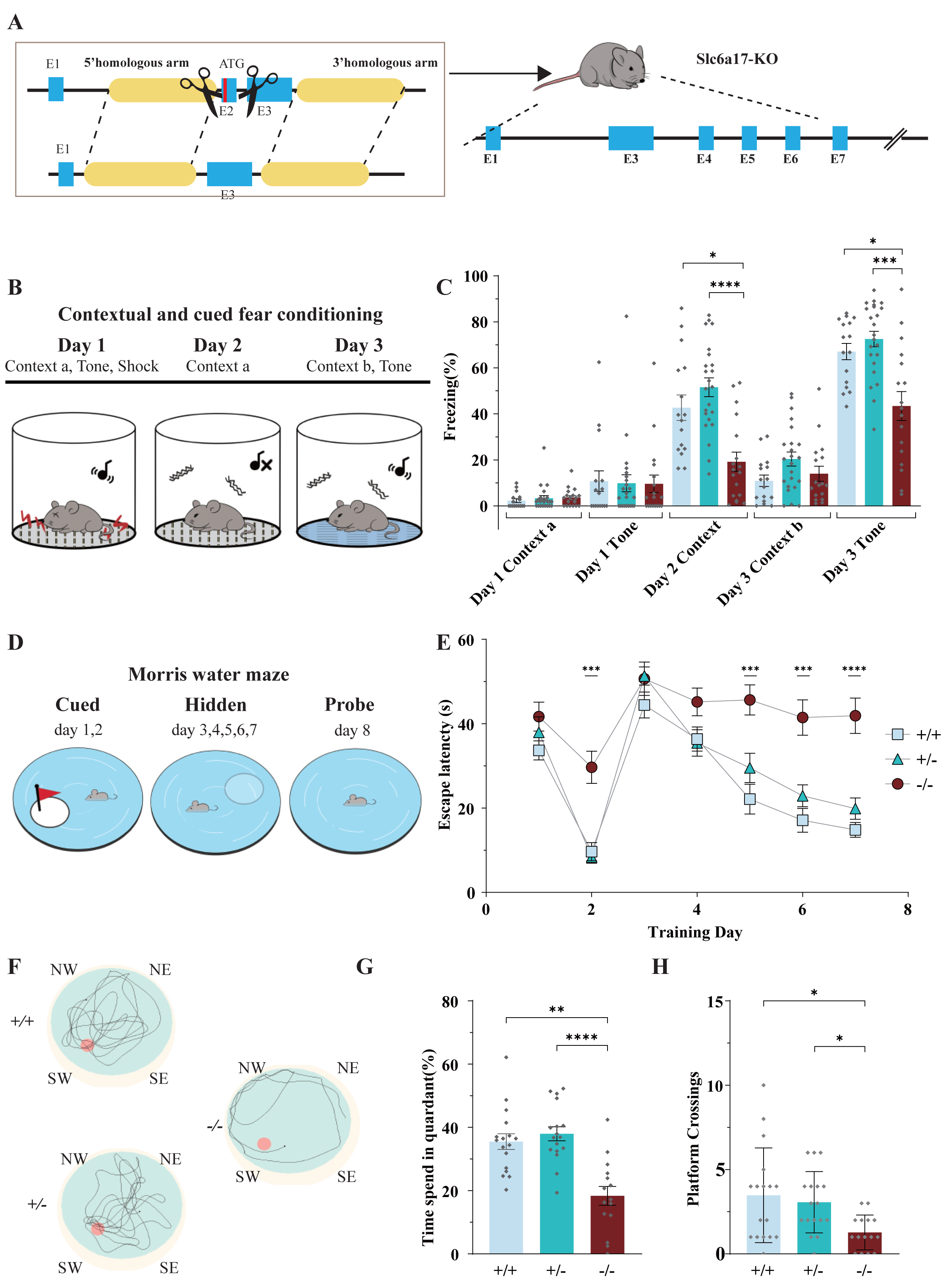
Slc6a17-KO Mice Exhibited Impaired Memory Formation. (A) A schematic diagram illustrating the knock-out strategy for generating Slc6a17-KO mice by CRISPR/Cas9. Exon 2 encoding the first 99 amino acids of Slc6a17 was deleted. More details in Figure S2A. (B) A diagram of experiments for contextual and cued fear conditioning (n=16, 23, 18 for Slc6a17^+/+^, Slc6a17^+/-^ and Slc6a17^-/-^, respectively). Detailed description in Experimental Procedures. (C) Slc6a17^-/-^ mice exhibited a significant decrease of freezing behavior in both cued and contextual fear conditioning (Day 2 context a, *p*=0.0062 for Slc6a17^+/+^ vs. Slc6a17^-/-^, *p*<0.0001 for Slc6a17^+/-^ vs. Slc6a17^-/-^; Day 3 tone, *p*=0.0093 for Slc6a17^+/+^ vs. Slc6a17^-/-^, *p*=0.0012 for Slc6a17^+/-^ vs. Slc6a17^-/-^). (D) A diagram of experiments for modified Morris water maze task (n=15, 18, 16 for Slc6a17^+/+^, Slc6a17^+/-^ and Slc6a17^-/-^, respectively). Detailed description in Experimental Procedures. (E) Latency to find the hidden platform during the training session. Slc6a17^-/-^ mice differred significantly from Slc6a17^+/+^ and Slc6a17^+/-^ mice (Two-way ANOVA; main effect of training day: F (4.681, 210.7) = 46.55, *p*<0.0001; main effect of genotype: F (2, 45) = 21.53, *p*<0.0001; main effect of training day x genotype: F (12, 270)=3.762, *p*< 0.0001; Tukey’s multiple comparisons test: day 2, *p*=0.0004 for Slc6a17^+/+^ vs. Slc6a17^-/-^, *p*=0.0002 for Slc6a17^+/-^ vs. Slc6a17^-/-^; day 5, *p*= 0.0002 for Slc6a17^+/+^ vs. Slc6a17^-/-^, *p*= 0.008 for Slc6a17^+/-^ vs. Slc6a17^-/-^; day 6, *p*=0.0002 for Slc6a17^+/+^ vs. Slc6a17^-/-^, *p*=0.0027 for Slc6a17^+/-^ vs. Slc6a17^-/-^; day 7, *p*<0.0001 for Slc6a17^+/+^ vs. Slc6a17^-/-^, *p*=0.0005 for Slc6a17^+/-^ vs. Slc6a17^-/-^). (F) Representative swimming traces of each genotype. (G) Percentage of time spent in the target quadrant was significantly different (*p*=0.0004 for Slc6a17^+/+^ vs. Slc6a17^-/-^, *p*< 0.0001 for Slc6a17^+/-^ vs. Slc6a17^-/-^). (H) Number of crossings through the platform area was significantly different (*p*=0.0194 for Slc6a17^+/+^ vs. Slc6a17^-/-^, *p*=0.0054 for Slc6a17^+/-^ vs. Slc6a17^-/-^). Here and hereafter, data in the Figure are presented as the mean ± SEM, with * indicating *p*< 0.05, ** indicating *p*< 0.01, *** indicating *p*< 0.001, and **** indicating *p*< 0.0001.

To investigate learning and memory of Slc6a17-KO mice, we first tested Slc6a17-KO mice on a classical associative learning model of contextual and cued fear (Hitti and Siegelbaum, 2014; Phillips and Ledoux, 1992) (Figure 2B). Both the WT and the heterozygous (Slc6a17^+/-^) mice spent a significant amount of time freezing (Figure 2C) on day 2 in Context-a environment. However, homozygous (Slc6a17^-/-^) mutants showed a dramatic decrease in freezing behavior compared to the other genotypes (Figure 2C). On day 3, both Slc6a17^+/+^ and Slc6a17^+/-^ mice showed significantly increased freezing behavior (Figure 2C), whereas Slc6a17^-/-^ mutants showed significant memory impairment in response to the tone (Figure 2C). These results indicate that the Slc6a17-KO mice were severely deficient in contextual and cued fear memory.

We then tested hippocampus-dependent spatial learning and memory by evaluating the performance of mice in the Morris water maze (Morris, 1984; Morris et al., 1982) (Figure 2D). During training, control (Slc6a17^+/+^ and Slc6a17^+/-^) mice showed a gradually decreasing latency to find the platform in both visible and hidden sessions (Figure 2E), whereas Slc6a17^-/-^ mutants exhibited significantly longer latency to find the platform (Figure 2E). Representative swimming traces of each genotype in probe trial are shown in Figure 2F. During the probe trial, control mice spent more time in the target quadrant (Figure 2F), whereas Slc6a17^-/-^ mutants spent much less time searching the platform quadrant (Figure 2G and 2H). The frequency that control animals crossed through the platform area was above chance level (Figure 2G and 2H), whereas Slc6a17^-/-^ mutants crossed the platform area much less than the other genotypes (Figure 2G and 2H).

Taken together, these data indicate that Slc6a17-KO mice are significantly impaired both in contextual and cued fear memory and in hippocampus-dependent spatial learning and memory.

### Behavioral deficits in Slc6a17^P663R^ mutant mice

We next generated a mouse line carrying a point mutation (P633R) found to be pathogenic in ID patients (Iqbal *et al*., 2015). Three repeats of the hemagglutinin (HA) epitope (Kolodziej and Young, 1991) and three repeats of the FLAG tag were fused in-frame to the C terminus of the endogenous P633R mutant (Figure 3A and S2B). Homozygous (Slc6a17^P633R/P633R^) mutant mice were not significantly different from the WT (Slc6a17^+/+^) or heterzygous (Slc6a17^P633R/+^) in body weight (Figure S4A), basal activities (Figure S4B) or novel objection recognition (Figure S4C).

**Figure 3.**
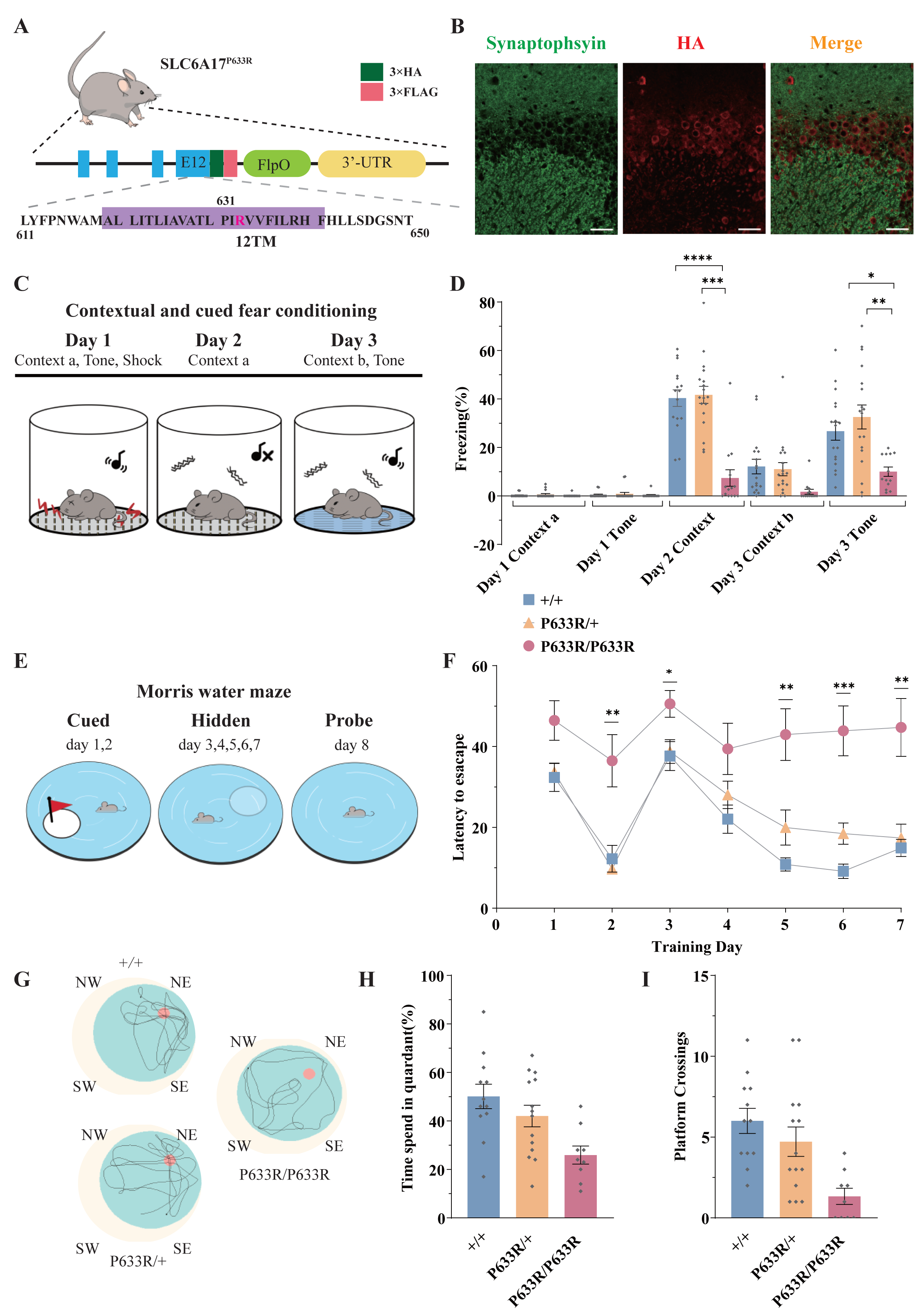
Deficient Memory in Slc6a17^P633R^ Mice. (A) A diagram illustrating the knock-in strategy for Slc6a17 with a pathogenic point mutation (Slc6a17^P633R^). It was tagged with 3 repeats of the HA epitope and 3 repeats of the FLAG epitope. More details in Figure S2B. (B) Immunocytochemistry of Slc6a17^P633R^ mouse brains with the anti-HA antibody showed that the subcellular localization of SLC6A17^P633R^-HA did not co-localized with synaptophysin. It was present in the cytoplasm. Representative view of CA3 was shown. Scale bar=50 μm. (C) Experiments for contextual and cued fear conditioning (n=17, 18, 14 for Slc6a17^+/+^, Slc6a17^P633R/+^, Slc6a17^P633R/P633R^, respectively). Detailed description in Experimental Procedures. (D) Slc6a17^P633R^ homozygous mutants exhibited significant decrease of freezing behavior in both cued and contextual fear conditioning (Day 2 context A, *p*< 0.0001 for Slc6a17^+/+^ vs. Slc6a17^P633R/P633R^, *p*< 0.0001 for Slc6a17^P633R/+^ vs. Slc6a17^P633R/P633R^; Day 3 tone, *p*= 0.0017 for Slc6a17^+/+^ vs. Slc6a17^P633R/P633R^, *p*= 0.0009 for Slc6a17^P633R/+^ vs. Slc6a17^P633R/P633R^). (E) Cartoon figure of experiments for Morris water maze task (n=12, 15, 9 for Slc6a17^+/+^, Slc6a17^P633R/+^ and Slc6a17^P633R/P633R^, respectively). Detailed description in Experimental Procedures. (F) Latency to find the hidden platform during the training session. Slc6a17^P633R/P633R^ mice showed significantly different learning curve from Slc6a17^+/+^ and Slc6a17^P633R/+^ mice (Two-way ANOVA; main effect of training day: F (4.381, 144.6) = 20.02, *p*<0.0001; main effect of genotype: F (2, 33) = 22.65, *p*<0.0001; main effect of training day x genotype: F (12, 198) = 2.389, *p*=0.0067; Tukey’s multiple comparisons test: day 2, *p*= 0.0150 for Slc6a17^+/+^ vs. Slc6a17^P633R/P633R^, *p*= 0.0078 for Slc6a17^P633R/+^ vs. Slc6a17^P633R/P633R^; day 3, *p*= 0.0403 for Slc6a17^+/+^ vs. Slc6a17^P633R/P633R^, *p*= 0.0388 for Slc6a17^P633R/+^ vs. Slc6a17^P633R/P633R^; day 5, *p*= 0.0023 for Slc6a17^+/+^ vs. Slc6a17^P633R/P633R^, *p*= 0.0243 for Slc6a17^P633R/+^ vs. Slc6a17^P633R/P633R^; day 6, *p*= 0.0199 for Slc6a17^+/+^ vs. Slc6a17^P633R/+^,*p*= 0.0010 for Slc6a17^+/+^ vs. Slc6a17^P633R/P633R^, *p*= 0.0077 for Slc6a17^P633R/+^ vs. Slc6a17^P633R/P633R^; day 7, *p*= 0.0074 for Slc6a17^+/+^ vs. Slc6a17^P633R/P633R^, *p*= 0.0129 for Slc6a17^P633R/+^ vs. Slc6a17^P633R/P633R^). (G) Representative swimming traces of each genotype. (H) Percentage of time spent in the target quadrant was significantly different (*p*= 0.0031 for Slc6a17^+/+^ vs. Slc6a17^P633R/P633R^, *p*= 0.0325 for Slc6a17^P633R/+^ vs. Slc6a17^P633R/P633R^). (I) Number of crossings through the platform area was significantly different (*p*= 0.0003 for Slc6a17^+/+^ vs. Slc6a17^P633R/P633R^, *p*= 0.0123 for Slc6a17^P633R/+^ vs. Slc6a17^P633R/P633R^). Data in the Figure are presented as the mean ± SEM, with * indicating *p*< 0.05, ** indicating *p*< 0.01, *** indicating *p*< 0.001, and **** indicating *p*< 0.0001.

We then investigated learning and memory in Slc6a17^P633R^ mice. In contextual and cued fear conditioning (Figure 3C), Slc6a17^P633R/P633R^ mutants showed significantly less freezing behavior compared to Slc6a17^+/+^ and heterozygous (Slc6a17^P633R/+^) mice on day 2 in Context-a environment (Figure 3D). On day 3, Slc6a17^P633R/P633R^ mutants showed a lower level of freezing behavior induced by cued tone compared to Slc6a17^+/+^ and Slc6a17^P633R/+^ mice (Figure 3D). These results indicate that Slc6a17^P633R^ mice are defective in contextual and cued fear memory.

We then tested Slc6a17^P633R^ mice in the Morris Water Maze for hippocampal spatial learning and memory (Figure 3E) (Morris, 1984; Morris *et al*., 1982). During training, Slc6a17^+/+^ and Slc6a17^P633R/+^ mice had a similar pattern in their learning curves with gradual decreases in the latency to find the platform in two sessions (Figure 3F). Similar to the Slc6a17-KO mice, Slc6a17^P633R/P633R^ mutants required significantly longer time to find the platform in both visible and hidden sessions (Figure 3F). Representative swimming traces of each genotype in probe trials are shown in Figure 3G. During the probe trial, Slc6a17^+/+^ and Slc6a17^P633R/+^ mice spent more time in the target quadrant (Figure 3H) and had more crossings through the platform area than Slc6a17^P633R/P633R^ mutants (Figure 3I).

Taken together, these data indicate that Slc6a17^P633R^ mice are also significantly impaired both in contextual and cued fear memory and in hippocampus-dependent long-term spatial learning. Because all the phenotypes of Slc6a17^P633R^ are similar to Slc6a7-KO, the point mutation P633 is a loss of function (LOF) mutation.

### Vesicular localization of SLC6A17

Previous studies with anti-SLC6A17 antibodies provided evidence that SLC6A17 was localized in SVs (Fischer *et al*., 1999; Kachidian, 1999; Masson *et al*., 1999). We have now used multiple approaches to analyze the subcellular localization of SLC6A17.

Slc6a17^-HA-2A-iCre^ mice were generated with the C terminus of the endogenous SLC6A17 protein fused in frame with three repeats of the HA epitope (Figure 4A and S1D). We used differential centrifugation to purify SVs from the brains of Slc6a17^-HA-2A-iCre^ mice (Evans, 2015; Huttner et al., 1983). As shown in Figure S5C, SLC6A17-HA was enriched in LP2 fraction, similar to the other SV proteins such as synaptotagmin 1 (Syt1) (Geppert et al., 1994), vesicular ATPase (V-ATPase) (Cidon and Sihra, 1989), and synaptobrevin 2 (Syb2) (Link et al., 1992; Schiavo et al., 1992), but different from postsynaptic proteins such as postsynaptic density 95 (PSD95) (Cho et al., 1992; Woods and Bryant, 1991). Sucrose gradient centrifugation of the LP2 fraction showed copurification of SLC6A17-HA with SV proteins including synaptophysin (Syp) (Jahn et al., 1985; Leube et al., 1987; Wiedenmann and Franke, 1985), Syt1, and VGluT1 (Bellocchio et al., 2000; Takamori et al., 2000), but not with proteins localized to the postsynaptic membrane (PSD95) or (SNAP23) (Ravichandran et al., 1996; Suh et al., 2010), the endoplasmic reticulum (ER) (ERp72), the endosome (EEA1) (Mu et al., 1995), the proteosome (PSMC6) or the trans-Golgi network (synaptaxin 6, STX6) (Figure 4D).

**Figure 4.**
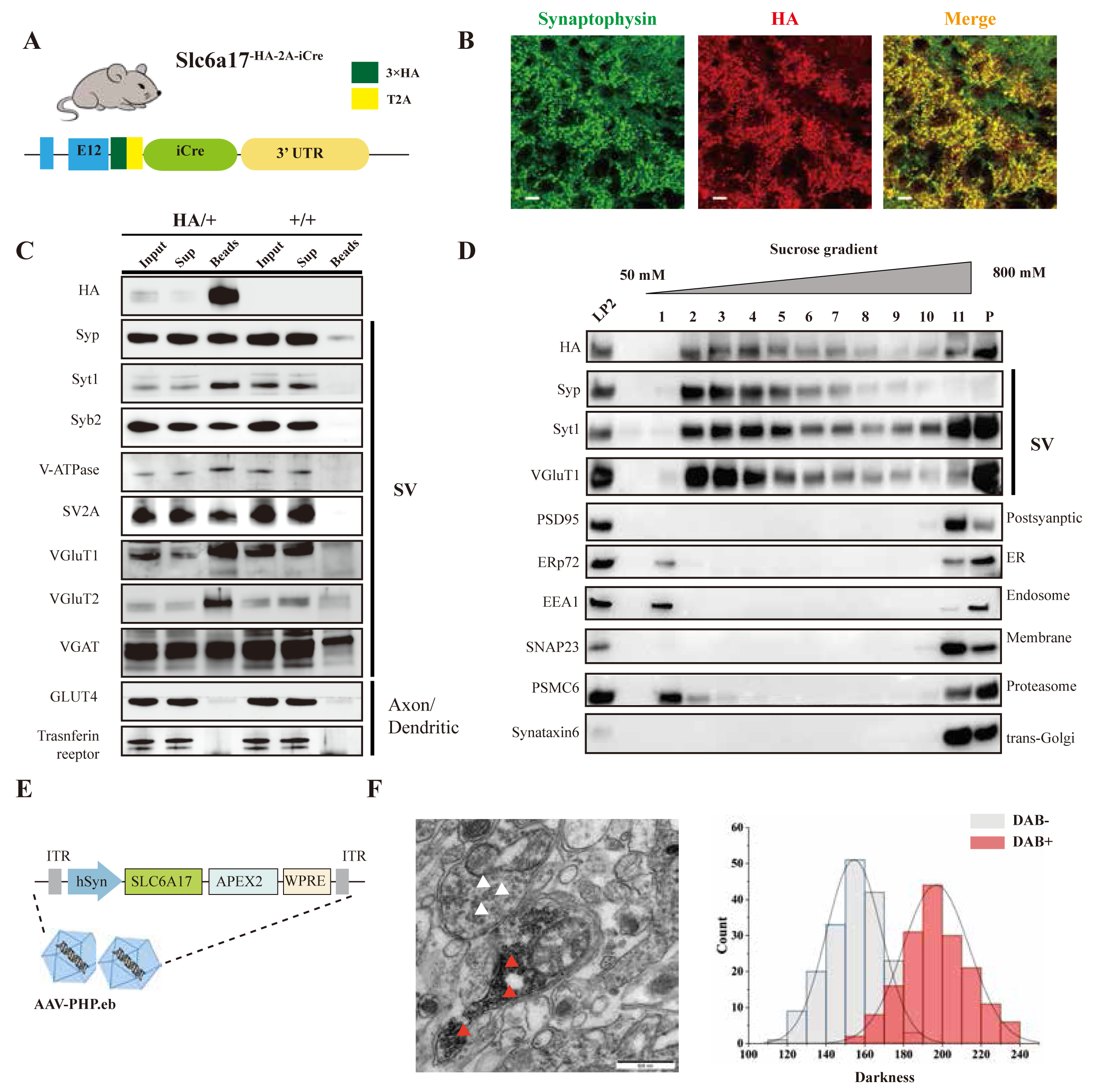
Biochemical and Genetically Assisted EM Validation of the Vesicular Localization of SLC6A17. (A) A schematic diagram illustrating the knock-in strategy for generating Slc6a17^-HA-2A-iCre^ mice. More details in Figure S1D. (B) Higher magnification views of co-immunostaining by anti-Syp and anti-HA antibodies in the hippocampus. Scale bar=10 μ . (C) Immunoisolation of SLC6A17-HA containing vesicles by the anti-HA antibody coated on magnetic beads. SLC6A17-HA fraction was positive for Syp, Syt1, Syb2, v-ATPase, VGluT1, VGluT2, and vGAT, but negative for GluT4 and transferin receptor. (D) Further purification of the LP2 fraction by sucrose gradient showed that SLC6A17-HA was co-immunoisolated with Syp, Syt1 and VGluT1, but not PSD95, ERp72, EEA1, SNAP23, PSMC6 or STX6. (E) A schematic diagram illustrating the APEX2-based labeling strategy with AAV-PHP.eb virus mediated hSLC6A17-APEX2 overexpression *in vivo*. SLC6A17 was fused in-frame to 3 repeats of the HA tag, a V5 tag, and APEX2. (F) Representative EM image of SVs labeled by hSLC6A17-APEX2 and darkness distribution of DAB positive and DAB negative SVs in sections of Slc6a17-APEX2 mouse brains. Red arrow pointing to APEX2 labeled SVs. White arrow pointing to unlabeled SVs.

We used the anti-HA antibody to immunoisolate SLC6A17-HA protein from Slc6a17^-HA-2A-iCre^ mice (Slc6a17^HA/+^), with WT (SLC6A17^+/+^) mice as a control (Figure 4C and S5D). After examining markers of subcellular organelles, the fraction containing SLC6A17-HA was found to be co-localized with SV markers such as Syp, Syt1, Syb2, VATPase and vesicular neurotransmitter transporters including VGluT1, VGluT2, and VGAT (Figure 4C and S5D), but not with markers of axons or dendrites (GLUT4 or transferrin receptor), ER (ERp72), the lysosome (LAMP2, LC3b or cathepsin D), the cytoplasmic membrane (SNAP23, PSD95 and GluN1), the Golgi apparatus (GM130 and Goglin-97), mitochondria (VADC), the active zone (ERC1b/2), the endosome (EEA1) and the proteosome (PSMC6) (Figure 4C and S5D). In addition, immunostaining of HA-tagged SLC6A17 in Slc6a17^HA/+^ mouse showed puncta co-localized with the anti-Syp positive immunoreactivity (Figure 4B, S2C and S5B).

We also used genetically assisted EM to confirm SLC6A17 localization. APEX2 is derived from ascorbate peroxidase (APEX) and can be used to genetically label proteins for EM inspection (Lam et al., 2015; Martell et al., 2017). We fused APEX2 in-frame to the C terminus of SLC6A17 (Figure 4E). SLC6A17-APEX2 fusion protein was expressed in the adult mouse brain by adeno-associated virus (AAV)-PHP.eb mediated transduction (Chan et al., 2017; Deverman et al., 2016) (Figure 4E). SLC6A17-APEX2 was indeed localized on the SVs (Figure 4F and S6A, S6C, S6E, S6G). Quantitative analysis showed distinct populations of SVs distinguished by electron density (Figure 4F and S6B, S6D, S6F, S6H).

Our results from biochemistry and genetically assisted EM analysis support the SV localization of SLC6A17.

### Functional significance of the SV localization of SLC6A17

We used a monoclonal anti-Syp antibody to immunoisolate SVs from brains of Slc6a17^+/+^ or Slc6a17^-/-^ mice (Boyken et al., 2013; Gronborg et al., 2010; Jahn *et al*., 1985). Quantitative proteomic analysis showed that only SLC6A17 was dramatically reduced in SVs immunopurified from Slc6a17-KO mice, whereas other transporters such as VGluT1, VGluT2, VGluT3, VGAT, VMAT2, SV2A, SV2B, SV2C, ZnT3 and VAT-1 were not different between Slc6a17^-/-^ and Slc6a17^+/+^ (Figure S7). Thus, Slc6a17 gene knockout indeed specifically decreased SLC6A17 protein, but none of the other transporters.

To investigate the functional significance of the subcellular localization of SLC6A17, we examined the subcellular localization of SLC6A17^P633R^ protein in the mouse brain. SLC6A17^P633R^ mutation was found in human ID patients (Iqbal et al., 2015) and transfection of a cDNA encoding this mutant into cultured primary neurons from mice found that its distribution was different from that of the WT SLC6A17 (Iqbal et al., 2015), but the disturbance on SV localization for the transfected mutant protein was not unambiguous and the effect of the mutation on the endogenous protein was unknown.

With the endogenous SLC6A17^P633R^ protein tagged by the HA and FLAG epitopes (Figure 3A and S2B), we found that SLC6A17^P633R^-HA signal was primarily localized to the soma with no overlap detected with the anti-Syp antibody staining (Figure 3B and S2C). We also performed differential centrifugation and sucrose gradient purification. HA and FLAG signals were presented in LP2 but did not co-purified with SVs in sucrose gradients, indicated that the SLC6A17^P633R^ protein was not present in SVs (Figure S2D).

### Presence of Gln in immunoisolated SVs

Neurotransmitters such as glutamate (Glu) and gamma-aminobutyric acid (GABA) have been reliably found in SVs immunoisolated from the mammalian brain (Bradberry et al., 2022; Burger et al., 1991; Burger et al., 1989; Chantranupong et al., 2020; Martineau et al., 2013). Previous experiments suggest that SLC6A17 could transport Ala, Gly, Leu and Pro into PC12 and CHO cells (Parra et al, 2008), or Ala, Gly, Leu, Pro, Cys, Gln, Gly, His and Ser into HEK cells (Zaia and Reimer, 2009). However, none of the reported substrates of SLC6A17 has been reliably detected in SVs.

To analyze the contents of SVs, we used the anti-Syp antibody to immunoisolate SVs from the mouse brain (Burger *et al*., 1991; Burger *et al*., 1989; Martineau *et al*., 2013), before their contents were subjected to chemical analysis by liquid chromatography coupled to high resolution mass spectrometry (LC-MS) (Figure 5A and 5B, and Figure S8B to S8G). IgG was used as a control for the anti-Syp antibody in SV immunoisolation. The specificity of the anti-Syp antibody mediated SVs immunoisolation was confirmed (Figure S8A, with a total of 16 markers for SVs, lysosomes, endosomes, the Golgi apparatus, mitochodria, the ER and pre-or post-synaptic membrane).

**Figure 5.**
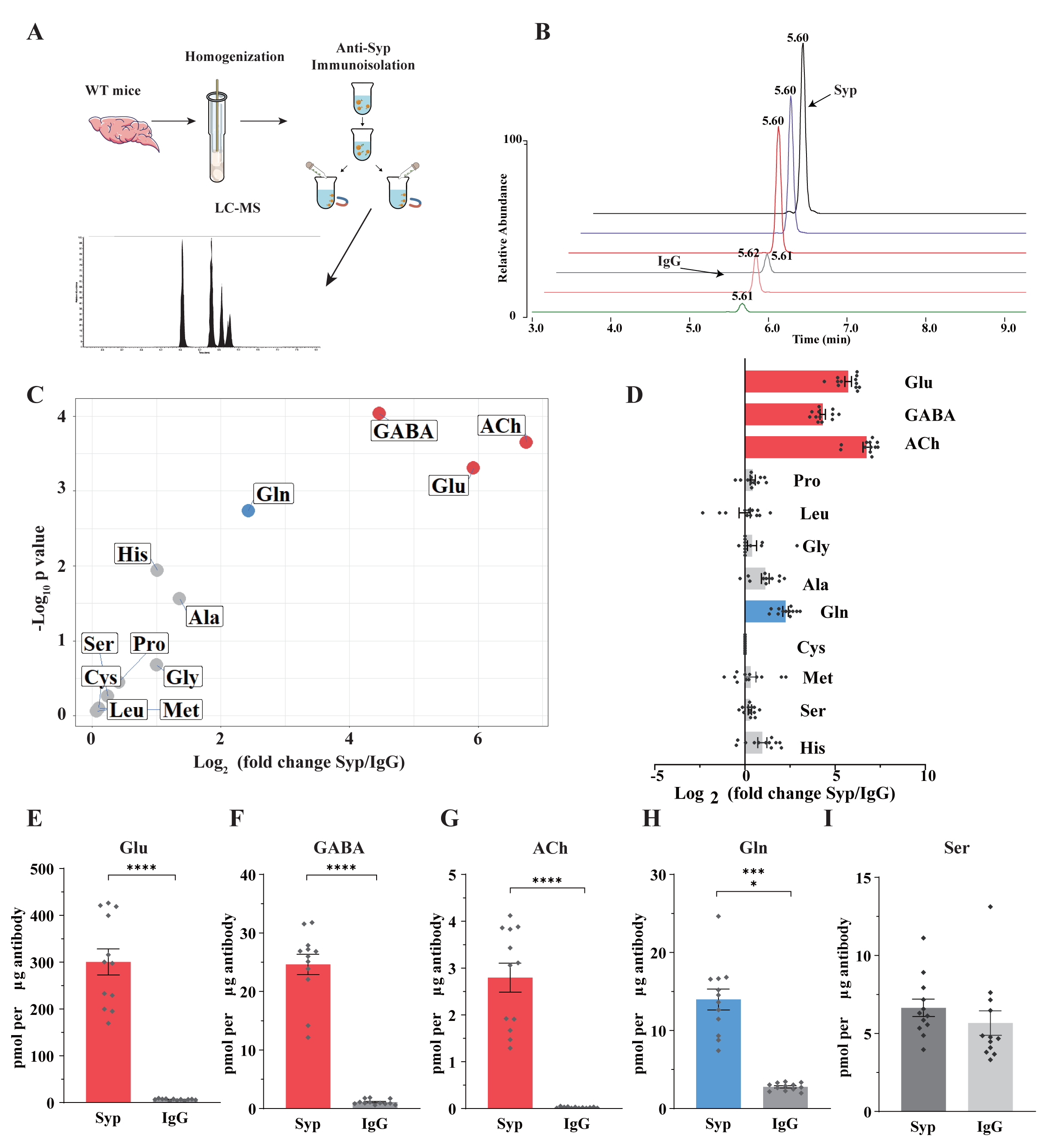
Presence of Glutamine in SVs from the Mouse Brain. **(A) A schematic** diagram illustrating the procedure to immunoisolate SVs with the anti-Syp antibody for LC-MS analysis of the contents in the SVs. (B) Representative MS results showing Gln signals from SVs immunoisolated by the anti-Syp antibody (anti-Syp) vs the control sample immunoisolated with IgG. (C) Volcano plot of chemical contents in the SVs isolated by anti-Syp vs IgG. The y axis shows *p* values in log_10_ and the x axis shows the log_2_ of the ratio of the level of a molecule immunoisolated by anti-Syp vs IgG. Classical neurotransmitters Glu, GABA and ACh, as well as previously reported substrates of SLC6A17 are listed. (D) Ratios of the level of a chemical immunoisolated by anti-Syp vs IgG (transformed into log_2_). (E-I) Chemicals were quantified to mole per μg antibody (n=12 for each group): Glu (E, *p*<0.0001 for anti-Syp vs. IgG); GABA (F, *p*<0.0001 for anti-Syp vs. IgG); ACh (G, *p*<0.0001 for anti-Syp vs. IgG); Gln (H, *p*<0.0001 for anti-Syp vs. IgG); Ser (I, *p*=0.7553 for anti-Syp vs. IgG).

The ratio of a molecule immunoisolated with the anti-Syp antibody to that with IgG was calculated and analyzed (Figure 5C and 5D). Glu, GABA and ACh were reliably detected from SVs immunoisolated with the anti-Syp antibody (Figure 5C to 5G, and S8B, S8D and S8F). Monoamine neurotransmitters such as 5-hydroxytryptamine (5-HT) (Figure S8C), dopamine (DA) (Figure S8G) and histamine (Figure S8E) could also be detected but not further analyzed for this paper. Hereafter, we focused our analysis on Glu, GABA and ACh as the positive controls and the nine previously reported AAs as candidate substrates of SLC6A17 *in vivo*.

Surprisingly, of the 9 AAs reported previously using cultured cells, 8 (Ala, Gly, Leu, Pro, Cys, Gly, His and Ser) were not found to be significantly enriched in SVs (Figure 5C and 5D). Only Gln was reproducibly found to be enriched in the anti-Syp antibody immunoisolated SVs (Figure 5C, 5D and 5H).

There could be several reasons why molecules transported into cultured cells were not detected in SVs of the mouse brain, including, e.g., relative abundance in subsets of SVs specific for different transporters, or the redundancy of multiple transporters for a single molecule among the 9 candidates. However, the positive finding of Gln in the SVs is definitive.

### Enrichment of Gln in SLC6A17 containing SVs from the mouse brain

The above results with SVs purified by the anti-Syp antibody revealed Gln presence in SVs, but did not show association of Gln with specific transporter on SV, because Syp is a universal marker of SVs (Jahn, Schiebler, Quimet and Greengard, 1985; Wiedenmann and Franke, 1985; Leube et al., 1987; Südhof et al., 1987). To determine the potential relationship of SLC6A17 with Gln in the SVs, we used magnetic beads coated with the anti-HA antibody to immunoisolate SLC6A17-positive SVs from Slc6a17^-HA-2A-iCre^ mice (Figure 4C and S5D).

The monoclonal anti-HA antibody could enrich SVs containing SLC6A17 from the brains of Slc6a17^-HA-2A-iCre^ mice but not from wt mice, as confirmed by the analysis of 23 markers (Figure 4C and S5D). SVs thus isolated (Figure 6A) were subject to chemical analysis with LC-MS (Figure 6A, 6B). Glu, GABA and ACh were detected in SVs containing SLC6A17-HA (Figure 6C to 6G), consistent with the immunoblot results that these SVs contained VGluT1, VGluT2 and VGAT (Figure 4C). Significant enrichment of Gln was detected in SVs containing SLC6A17-HA (Figure 6C, 6D and 6H). By contrast, the other 8 AAs had not been found to be enriched in SVs containing SLC6A17-HA (Figure 6C, 6D and 6I).

**Figure 6.**
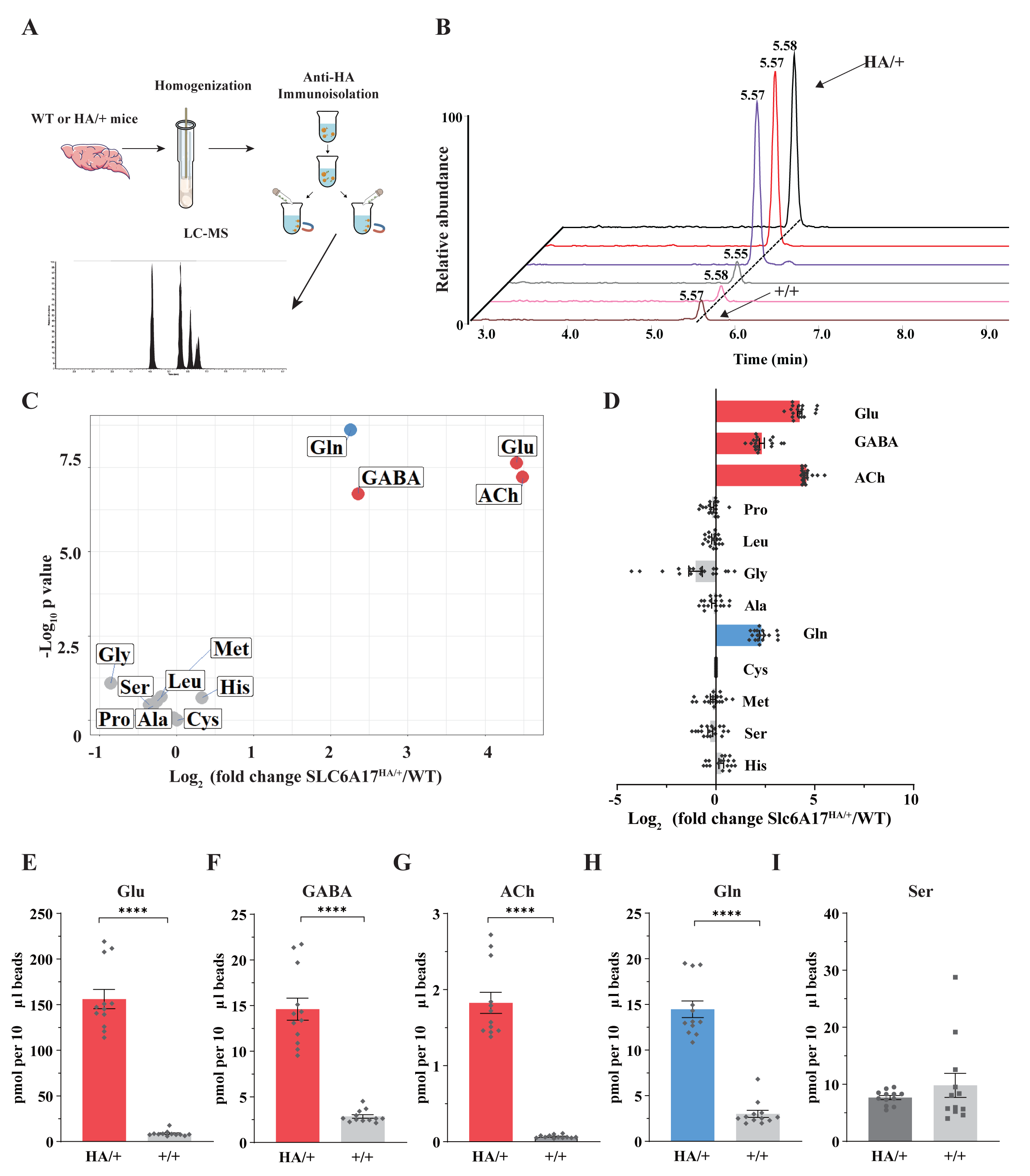
Gln Enrichment in SVs Containing SLC6A17. (A) A schematic diagram illustrating the procedure to isolate SLC6A17-containing SVs for chemical analysis of SV contents. (B) Representative result showing MS Gln signals in SVs immunoisolated by the anti-HA beads in SLC6A17^HA/+^ mice vs those from Slc6a17^+/+^ mice. (C) Volcano plot of chemical contents in the SVs immunoisolated by anti-HA from Slc6a17^HA/+^ mice vs Slc6a17^+/+^ mice. The y axis shows *p* values in log_10_ and the x axis shows the log_2_ of the ratio of the level of a molecule immunoisolated by anti-HA from Slc6a17^HA/+^ mice vs that from Slc6a17^+/+^ mice. Classical neurotransmitters such as Glu, GABA and ACh as well as the previously reported substrates of SLC6A17 are listed. (D) Ratios of the level of a chemical immunoisolated by anti-HA from Slc6a17^HA/+^ mice vs that from Slc6a17^+/+^ mice (transformed into log_2_). (E-I) Contents of SLC6A17-containing SVs were quantified to mole per 10 μl anti-HA beads (n=12, for each group): Glu (E, *p*<0.0001 for Slc6a17^HA/+^ vs. Slc6a17^+/+^); GABA (F, *p*<0.0001 for Slc6a17^HA/+^ vs. Slc6a17^+/+^); ACh (G, *p*<0.0001 for Slc6a17^HA/+^ vs. Slc6a17^+/+^); Gln (H, *p*< 0.0001 for Slc6a17^HA/+^ vs. Slc6a17^+/+^); Ser (I, *p*= 0.7553 for Slc6a17^HA/+^ vs. Slc6a17^+/+^).

Thus, only Gln has been reproducibly found to be enriched in SLC6A17 containing SVs *in vivo*.

### Sufficiency of SLC6A17 for Gln localization in SVs

To further investigate whether SLC6A17 could increase Gln in SVs, we overexpressed SLC6A17 in the mouse brain. AAV-PHP.eb was used to express hSLC6A17-HA in the mouse brain (OE-hSLC6A17-HA) (Chan *et al*., 2017; Deverman *et al*., 2016) (Figure 7A). The overexpressed hSLC6A17-HA protein was localized in SVs, as analyzed by immunoblot (Figure S9A).

**Figure 7.**
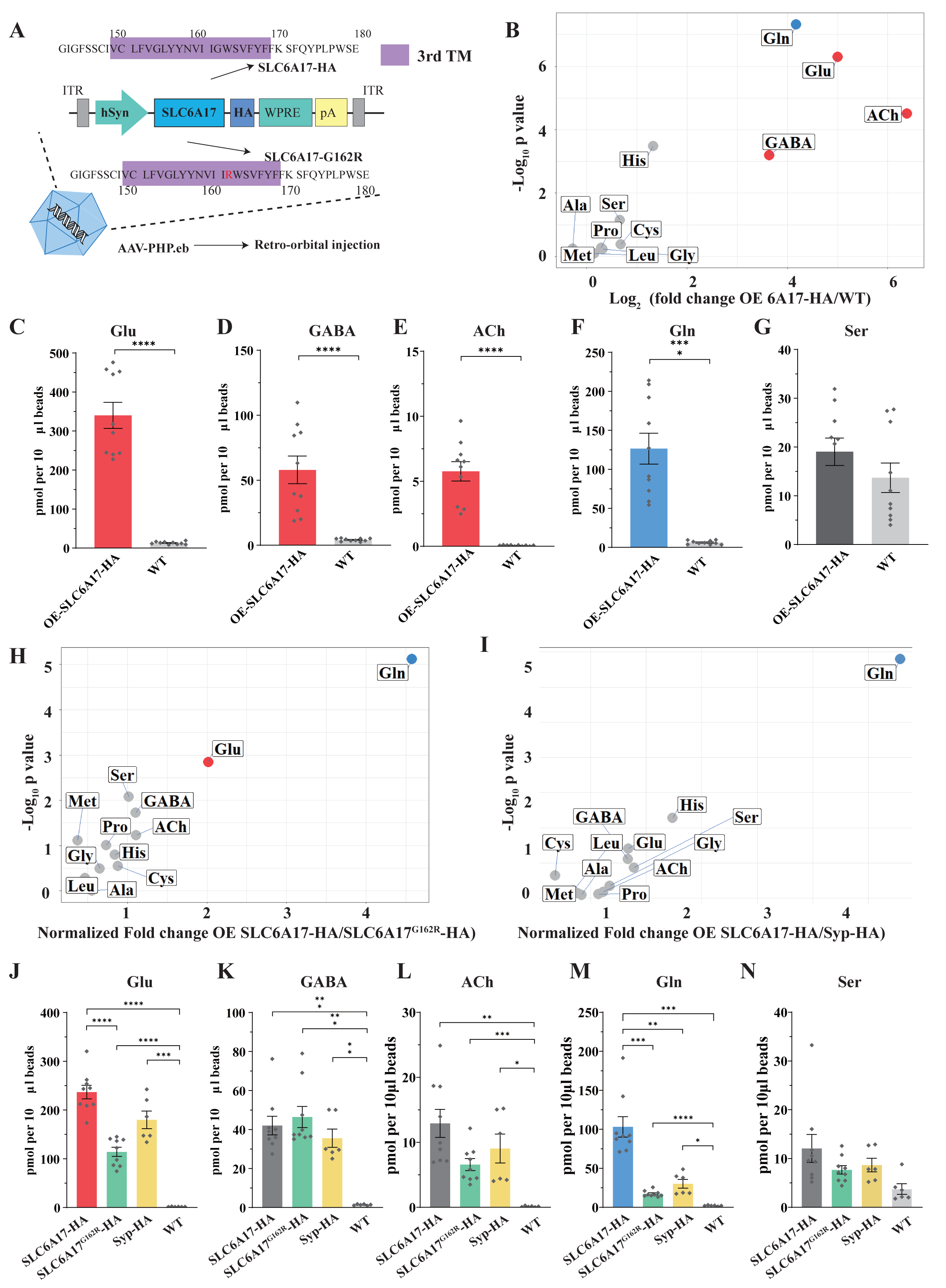
Increased Gln Transport into SVs Containing SLC6A17 but not into SVs Containing SLC6A17^G162R^. (A) A schematic diagram illustrating the strategy for the AAV-PHP.eb virus mediated *in vivo* overexpression of hSLC6A17-HA and hSLC6A17^G162R^-HA. (B) Volcano plot of chemical contents in the SVs isolated by anti-HA from the OE-hSLC6A17-HA mice vs WT mice. The y axis shows *p* values in log_10_ and the x axis shows the log_2_ of the ratio of the level of a molecule immunoisolated by anti-HA beads from mice overexpressing hSLC6A17-HA vs that from WT mice. Classical neurotransmitters and previously reported substrates of SLC6A17 are listed. (C-G) Contents of OE-hSLC6A17-containing SV are quantified to mole per 10 μl HA beads (n=10, for each group): Glu (C, *p*<0.0001 for OE-hSLC6A17-HA vs. WT); GABA (D, *p*<0.0001 for OE-hSLC6A17-HA vs. WT); ACh (E, *p*<0.0001 for OE-hSLC6A17-HA vs. WT); Gln (F, *p*<0.0001 for OE-hSLC6A17-HA vs. WT); (G, *p*=0.1655 for OE-hSLC6A17-HA vs. WT). (H) Volcano plot comparing the chemical contents of SVs containing OE-hSLC6A17-HA with those containing OE-hSLC6A17^G162R^-HA. Classical neurotransmitters and 9 putative substrates of SLC6A17 are listed. Glu is the only classical transmitter significantly increased. Gln is the only substrates significantly increased in hSLC6A17-HA containing SVs vs hSLC6A17^G162R^-HA containing SVs. (I) Volcano plot comparing the chemical contents of SVs containing OE-hSLC6A17-HA with those containing OE-Syp-HA. Classical neurotransmitters and 9 putative substrates of SLC6A17 are listed. Gln is the only substrates significantly increased in SLC6A17 containing SVs vs Syp-HA containing SVs. (J-N) Contents of SVs containing SCL6A17-HA, SVs containing OE-SLC6A17^G162R^-HA, SVs containing Syp-HA immunosilated by anti-HA immunoisolation with that from WT mouse brains were quantified to mole per 10 μl HA beads (n=9, 9, 6, 6, for OE-hSLC6A17-HA, OE-hSLC6A17^G162R^-HA, OE-Syp-HA, and WT, respectively): Glu (J, *p*<0.0001 for OE-hSLC6A17-HA vs. OE-hSLC6A17^G162R^-HA; *p*<0.0001 for OE-hSLC6A17-HA vs. WT; *p*<0.0001 for OE-hSLC6A17^G162R^-HA vs. WT; *p*= 0.0009 for OE-Syp-HA vs. WT; *p*= 0.1551 for OE-hSLC6A17-HA vs. OE-Syp-HA; *p*= 0.0618 for OE-hSLC6A17^G162R^-HA vs. OE-Syp-HA); GABA (K, *p*= 0.0002 for OE-hSLC6A17-HA vs. WT; *p*= 0.0002 for OE-hSLC6A17^G162R^-HA vs. WT; *p*= 0.0039 for OE-Syp-HA vs. WT; *p*= 0.9885 for OE-hSLC6A17-HA vs. OE-hSLC6A17^G162R^-HA; *p*= 0.9000 for OE-hSLC6A17-HA vs. OE-Syp-HA; *p*= 0.5840 for OE-hSLC6A17^G162R^-HA vs. OE-Syp-HA); ACh (L *p*= 0.0019 for OE-hSLC6A17-HA vs. WT; *p*= 0.0006 for OE-hSLC6A17^G162R^-HA vs. WT; *p*= 0.0467 for OE-Syp-HA vs. WT; *p*= 0.1054 for OE-hSLC6A17-HA vs. OE-hSLC6A17^G162R^-HA; *p*= 0.7563 for OE-hSLC6A17-HA vs. OE-Syp-HA; *p*= 0.8702 for OE-hSLC6A17^G162R^-HA vs. OE-Syp-HA); Gln (M, *p*= 0.001 for OE-hSLC6A17-HA vs. OE-hSLC6A17^G162R^-HA; *p*= 0.0018 for OE-hSLC6A17-HA vs. OE-Syp-HA; *p*= 0.0003 for OE-hSLC6A17-HA vs. WT; P<0.0001 for OE-hSLC6A17^G162R^-HA vs. WT; *p*= 0.0189 for OE-Syp-HA vs. WT; *p*= 0.2749 for OE-hSLC6A17^G162R^-HA vs. OE-Syp-HA); Ser (N, *p*= 0.626 for OE-hSLC6A17-HA vs. OE-hSLC6A17^G162R^-HA; *p*= 0.8551 for OE-hSLC6A17-HA vs. OE-Syp-HA; *p*= 0.9874 for OE-hSLC6A17-HA vs. OE-Syp-HA; P<0.0865 for OE-hSLC6A17^G162R^-HA vs. WT; *p*= 0.1017 for OE-Syp-HA vs. WT).

We then analyzed the chemical contents of immunoisolated SVs by LC-MS. The ratio of a molecule purified from OE-hSLC6A17-HA mice by the anti-HA antibody vs that purified from WT mice by the same antibody was calculated and analyzed (Figure 7B). SVs from mice overexpressing hSLC6A17-HA contained significantly higher levels of Glu and GABA (Figure 7B to 7G). Gln was dramatically increased (Figure 7B and 7F), to the extent that Gln was higher than GABA in SVs overexpressing hSLC6A17-HA (Figure 7B, 7D, 7F, 7K and 7M). Quantification of the enriched molecules in SVs containing hSLC6A17-HA showed Gln to be approximately 2 times that of GABA (Figure 7D and 7F). In addition, His was moderately increased in SVs overexpressing hSLC6A17-HA (Figure 7B, S9C and S9D). These results from our virally introduced overexpression experiments support that SLC6A17 is sufficient for Gln transport into SVs *in vivo*.

### Functional significance of SLC6A17 for Gln presence in the SVs

SLC6A17^G162R^ is another pathogenic mutation of human ID (Iqbal *et al*., 2015), with a point mutation in the 3^rd^ transmembrane domain of SLC6A17. To investigate whether SLC6A17^G162R^ affected Gln transportation, we used AAV-PHP.eb to overexpress either hSLC6A17^G162R^-HA or Syp-HA in mouse brains. Overexpressed hSLC6A17^G162R^-HA was still localized on SVs (Figure S9A and S9E). Overexpressed Syp-HA was previously reported to be localized on SVs (Chantranupong *et al*., 2020), which we confirmed (Figure S9A).

LC-MS analysis of contents of SVs immunoisolated by the anti-HA antibody from OE-Syp-HA mice, when compared with immunoprecipitates by the anti-HA antibody from the brains of WT mice, would indicate SV enriched molecules. As expected, Glu, GABA and ACh were all found to be enriched in these SVs (Figure 7J, 7K, and 7L), whereas Ser was not enriched (Figure 7N).

The contents of SVs immunoisolated from OE-hSLC6A17^G162R^-HA mice, when compared with that from OE-Syp-HA and WT mice, would indicate molecules enriched in SVs containing hSLC6A17^G162R^ mutation, above the general contents of all SVs. Levels of Glu and GABA in SVs containing hSLC6A17^G162R^-HA were higher than those in WT mice (Figure 7J, 7K and S9B), but not significantly different with those from OE-Syp-HA mice (Figure 7J and 7K). These results indicate that the subset of SVs containing hSLC6A17^G162R^-HA did not enrich Glu or GABA beyond the levels of Glu and GABA in the general population of SVs. The level of Gln in SVs containing hSLC6A17^G162R^-HA were much lower than that in from OE-hSLC6A17-HA mice (Figure 7M), indicating that hSLC6A17^G162R^ was defective in transporting Gln into SVs *in vivo*. The level of Gln in SVs containing hSLC6A17^G162R^-HA was higher than that in WT mice, but not different from that in OE-Syp-HA mice (Figure 7M). These results indicate that Gln was present in SVs, but not enriched in SVs containing SLC6A17^G162R^-HA over the general populations of SVs expressing Syp-HA. The levels of Gln and the other 8 AAs in SVs containing hSLC6A17^G162R^-HA were not significantly higher than those in WT mice (Figure 7N, S9B and S9D).

We compared the contents of SVs immunoisolated from OE-hSLC6A17-HA mice with those from either OE-Syp-HA mice or OE-SLC6A17^G162R^-HA mice. The levels of GABA and ACh were not different among SVs from mice expressing SLC6A17-HA, Syp-HA, or SLC6A17^G162R^-HA (Figure 7I, 7I, 7K and 7L), indicating that SVs containing SLC6A17-HA were similar to the general populations of SVs. A moderate enrichment of Glu in SVs overexpressing SLC6A17-HA, as compared to those overexpressing SLC6A17^G162R^-HA, was observed (Figure 7H and 7J). Gln was increased most dramatically in SVs overexpressing SLC6A17-HA, as compared to those overexpressing Syp-HA (Figure 7I and 7M) or SLC6A17^G162R^-HA (Figure 7H and 7M). The levels of Ser (Figure 7N), Pro, Gly, Met, Leu, Ala, Cys were not significantly increased in SVs overexpressing SLC6A17-HA, as compared to those overexpressing Syp-HA (Figure 7I) or SLC6A17^G162R^-HA (Figure 7H).

Taken together, our data provide *in vivo* evidence that the pathogenic mutation SLC6A17^G162R^ could not transport Gln into SVs in the mouse brain, supporting the functional significance of Gln in the SVs.

### Requirement of SLC6A17 for Gln transport into the SV in Slc6a17 knockout mice

We have generated SLC6A17-KO mice (Figure 2A) and found them to be defective in learning and memory (Figure 2B to 2H). We then investigated Gln level in these mice. The anti-Syp antibody was used to immunoisolate SVs from the brains of SLC6A17^+/+^, SLC6A17^+/-^ and SLC6A17^-/-^ mice (Figure 8A-8L).

**Figure 8.**
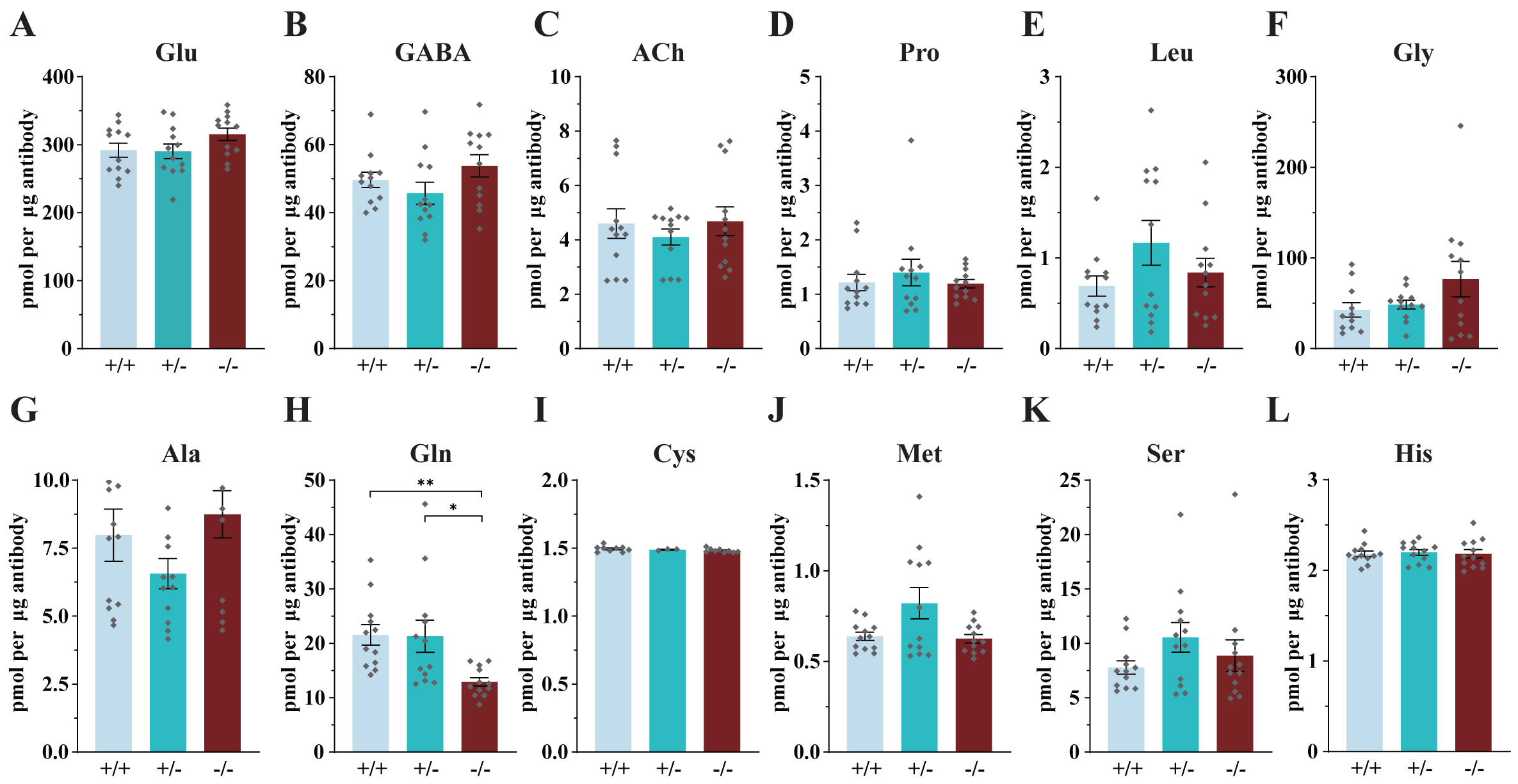
Reduced Levels of Gln in the SVs of Slc6a17 KO Mice. Contents of SVs purified by the anti-Syp antibody from Slc6a17-KO mice were quantified to mole per 10 μg antibody (n=12 for Slc6a17, Slc6a17, Slc6a17): Glu (A, *p*=0.2749 for Slc6a17^+/+^ vs. Slc6a17^-/-^, *p*=0.2466 for Slc6a17^+/-^ vs. Slc6a17^-/-^); GABA (B, *p*=0.665 for Slc6a17^+/+^ vs. Slc6a17^-/-^, *p*=0.2503 for Slc6a17^+/-^ vs. Slc6a17^-/-^); ACh (C, *p*=0.9993 for Slc6a17^+/+^ vs. Slc6a17^-/-^, *p*=0.7186 for Slc6a17^+/-^ vs. Slc6a17^-/-^); Pro (D, *p*=0.9986 for Slc6a17^+/+^ vs. Slc6a17^-/-^, *p*=0.8035 for Slc6a17^+/-^ vs. Slc6a17^-/-^); Leu (E, *p*=0.826 for Slc6a17^+/+^ vs. Slc6a17^-/-^, *p*= 0.61 for Slc6a17^+/-^ vs. Slc6a17^-/-^); Gly (F, *p*= 0.3211 for Slc6a17^+/+^ vs. Slc6a17^-/-^, *p*=0.442 for Slc6a17^+/-^ vs. Slc6a17^-/-^); Ala (G, *p*=0.9092 for Slc6a17^+/+^ vs. Slc6a17^-/-^, *p*=0.1321 for Slc6a17^+/-^ vs. Slc6a17^-/-^); Gln (H, *p*=0.002 for Slc6a17^+/+^ vs. Slc6a17^-/-^, *p*=0.0489 for Slc6a17^+/-^ vs. Slc6a17^-/-^); Cys (I, *p*=0.3753 for Slc6a17^+/+^ vs. Slc6a17^-/-^, *p*=0.5718 for Slc6a17^+/-^ vs. Slc6a17^-/-^); Met (J, *p*=0.9735 for Slc6a17^+/+^ vs. Slc6a17^-/-^, *p*= 0.13 for +/- vs. Slc6a17^-/-^); Ser (K, *p*=0.8678 for Slc6a17^+/+^ vs. Slc6a17^-/-^, *p*=0.7834 for Slc6a17^+/-^ vs. Slc6a17^-/-^); His (L, *p*>0.9999 for Slc6a17^+/+^ vs. Slc6a17^-/-^, *p*=0.9922 for Slc6a17^+/-^ vs. Slc6a17^-/-^).

LC-MS analysis of SVs revealed that Gln was the only molecule found to be decreased in SVs from Slc6a17^-/-^ mice, as compared with Slc6a17^+/+^ or Slc6a17^+/-^ mice (Figure 8H). Quantitative analysis showed that the levels of classic transmitters such as Glu (Figure 8A), GABA (Figure 8B) and ACh (Figure 8C) were not significantly different among the SVs immunoisolated from the Slc6a17^+/+^, the Slc6a17^+/-^ or the Slc6a17^-/-^ mice. The levels of the other 8 AAs were also not significantly different among SVs immunoisolated from all 3 genotypes (Figure 8D, 8E, 8F, 8G, 8I, 8J, 8K and 8L). These results further support that Gln was transported into the SVs by SLC6A17 *in vivo*.

### Physiological requirement of SLC6A17 for Gln presence in the SVs in adult mice

To investigate whether the endogenous SLC6A17 is required in adulthood for Gln presence in the SVs, we used a viral mediated CRISPR-based approach to remove the Slc6a17 gene from adult mice.

Briefly, Slc6a17^-HA-2AiCre^ mice generated earlier (Figure S1D) were crossed with LSL-Cas9 transgene-carrying mice (Platt et al., 2014) and injected with an AAV-PHP.eB virus to express Syp-HA and guide RNAs (gRNA) targeting Slc6a17 (AAV-PHP.eb-hSyn-DIO-Syp-HA-U6-sgRNAs) (Figure 9A and S10A). A tandemly arrayed tRNA-gRNA structure was used to maximize the cleavage possibility *in vivo* (Port and Bullock, 2016; Xie et al., 2015) (Figure S9A). This strategy caused the excision of the Slc6a17 gene and simultaneous expression of Syp-HA to tag SVs specifically in Slc6a17-expressing cells of adult mice (Figure S10A). Slc6a17-HA was significantly reduced both in the homogenates of total brains and in immunoisolated SVs from AAV-sgRNA/Slc6a17^iCre^/Cas9^+^ mice (Figure 9B and S9B).

**Figure 9.**
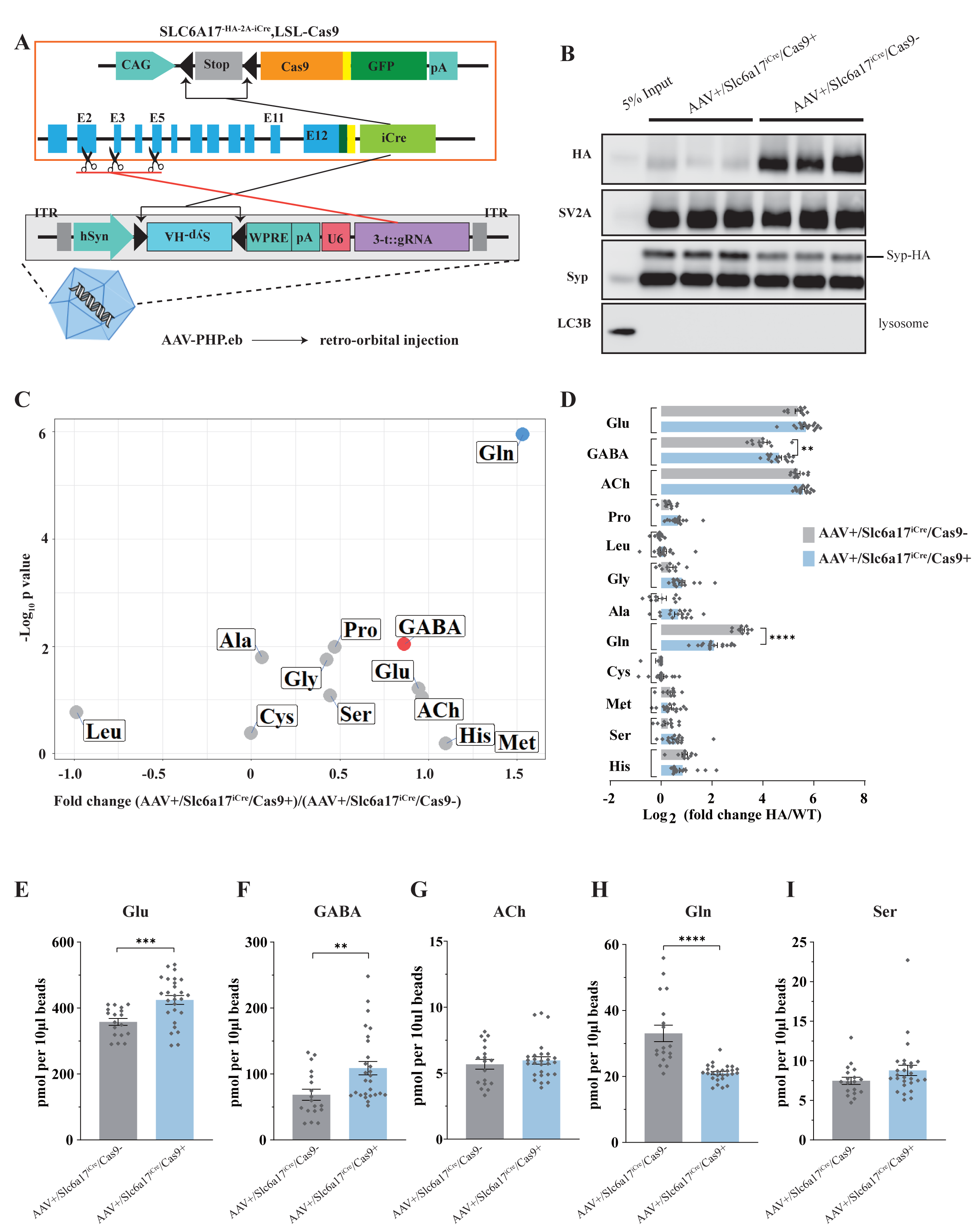
Physiological Requirement of SLC6A17 for Gln Transport into SVs in vivo. (A) A schematic diagram illustrating the strategy of Cas9-mediated cleavage of Slc6a17 specifically in Slc6a17 positive neurons, and simultaneous labeling of all SVs in these neurons by Syp-HA. (B) Immunoblot results showing SLC6A17 protein was significantly reduced in targeted neurons, while Syp-HA was efficiently tagged onto the SVs of these neurons. (C) Volcano plot of contents of SVs from AAV-sgRNA/Slc6a17^iCre^/Cas9^+^ targeted neurons compared to contents of SVs from control (AAV-sgRNA /Slc6a17^iCre^/Cas9^-^) neurons. Glu, GABA, ACh and the 9 previously reported substrates of SLC6A17 are listed. (D) Ratios of the level of a molecule from the SVs of AAV-sgRNA /Slc6a17^iCre^/Cas9^+^ neurons vs the level of the same molecule from SVs of AAV-sgRNA /Slc6a17^iCre^/Cas9^-^ neurons shown as fold change (log2 transformed). GABA level was significantly increased (*p*=0.0034 for AAV-sgRNA/Slc6a17^iCre^/Cas9^+^ vs. AAV-sgRNA /Slc6a17^iCre^/Cas9^-^). Gln level was significantly decreased (*p*< 0.0001 for AAV-sgRNA /Slc6a17^iCre^/Cas9^+^ vs. AAV-sgRNA/Slc6a17^iCre^/Cas9^-^). (E-H) Contents of SVs from Slc6a17 containing neurons were quantified to mole per 10 μl HA beads (n=18, 27 for Slc6a17 /Cas9 and Slc6a17^iCre^/Cas9^+^, respectively): Glu (E, *p*= 0.0005 for AAV-sgRNA/Slc6a17^iCre^/Cas9^+^ vs. AAV-sgRNA /Slc6a17^iCre^/Cas9^-^); GABA (F, *p*=0.0032 for AAV-sgRNA /Slc6a17^iCre^/Cas9^+^ vs. AAV-sgRNA /Slc6a17^iCre^/Cas9^-^); Gln (G, *p*<0.0001 for AAV-sgRNA /Slc6a17^iCre^/Cas9^+^ vs. AAV-sgRNA /Slc6a17^iCre^/Cas9^-^); Ser (H, *p*=0.0979 for AAV-sgRNA /Slc6a17^iCre^/Cas9^+^ vs. AAV-sgRNA/Slc6a17^iCre^/Cas9^-^).

Importantly, when the SV contents from the brains of knockout (AAV-sgRNA/Slc6a17^iCre^/Cas9^+^) and control (AAV-sgRNA/Slc6a17^iCre^/Cas9^-^) mice were compared after LC-MS analysis, Gln was the only vesicular content significantly decreased in SVs from the knockout mice (Figure 9C, 9D and 9H).

To our surprise, GABA and Glu were moderately increased (Figure 9C, 9D, 9E and 9F). The level of ACh, and those of the other 8 AAs (Ala, Gly, Leu, Pro, Cys, Gly, His and Ser) were not significantly changed in the SVs from adult Slc6a17 knockout mice (Figure 9C, 9D, 9G and 9I). The increased levels of Glu and GABA in SVs of virally mediated knock-out mice were different from the absence of Glu and GABA increases in the straightforward Slc6a17 KO (Figure 8A, 8B), making the significance of the observed increases of Glu and GABA in the virally mediated Slc6a17 knockout unclear.

These results indicate that SLC6A17 is physiologically necessary for Gln in SVs, but not for the other 8 AAs.

### Functional significance of Gln in SVs supported by analysis of mice carrying a mutation mimicking a human ID mutation

To further address the question whether Gln transport into the SVs was functionally important, we examined the SVs from Slc6a17^P633R^ mutant mice (Figure 10A to 10L). This mutation mimicked one of the human ID mutations (Iqbal et al., 2015), and its behavioral phenotypes have been found by us (Figure 3). We have shown that the SLC6A17^P633R^ protein was not localized in the SVs (Figure 3B and S2D).

**Figure 10.**
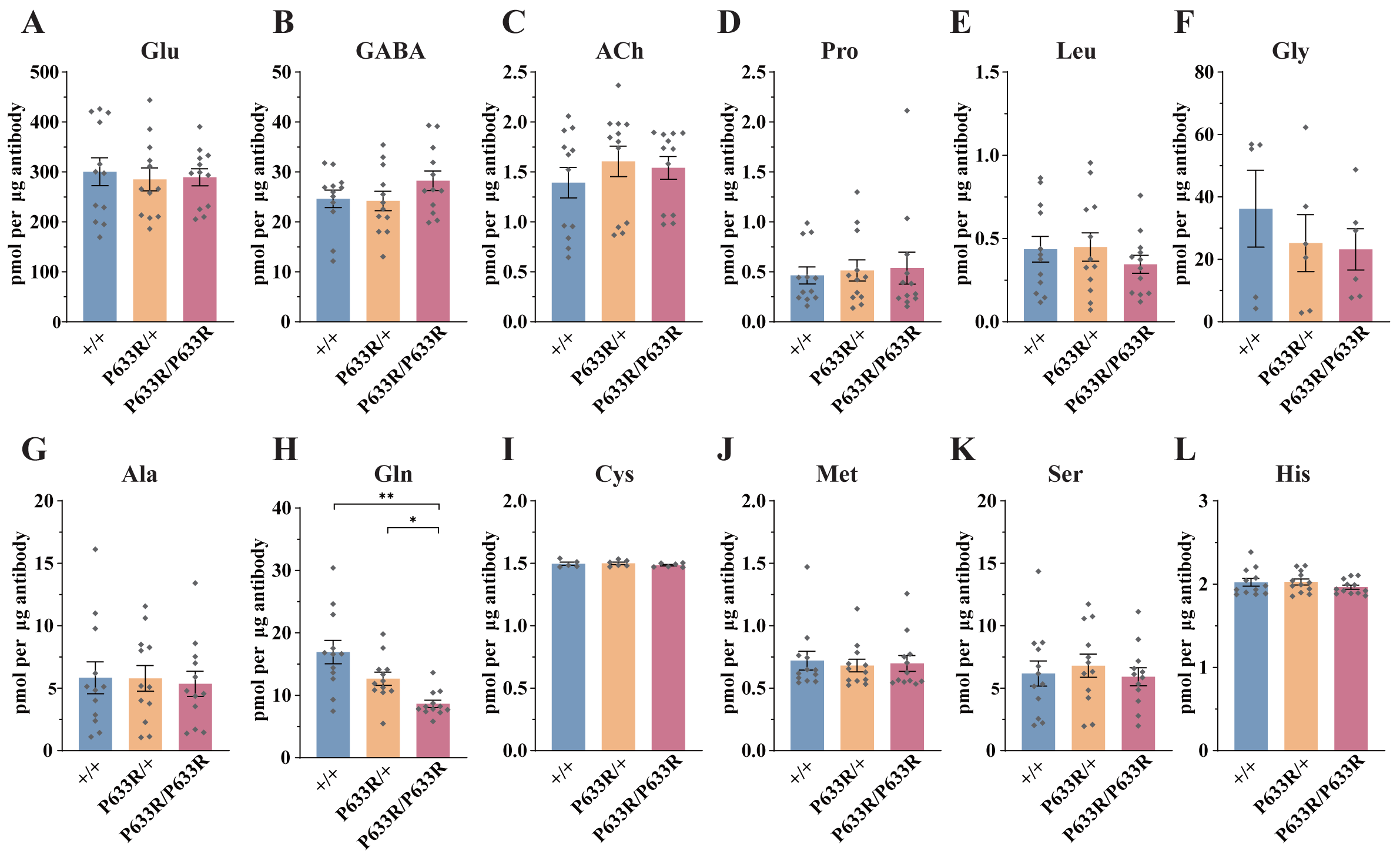
Reduced Levels of Gln in the SVs of Slc6a17^P633R^ Mice. Contents of SVs purified by the anti-Syp antibody from Slc6a17^P633R^ mice were quantified to mole per 10 μ antibody (n=12 for all genotype): Glu (A, *p*=0.9804 for Slc6a17 vs. Slc6a17^P633R/P633R^, *p*=0.9982 for Slc6a17^P633R/+^ vs. Slc6a17^P633R/P633R^); GABA (B, *p*=0.4432 for Slc6a17^+/+^ vs. Slc6a17^P633R/P633R^ *p*=0.3898 for Slc6a17^P633R/+^ vs. Slc6a17^P633R/P633R^); ACh (C, *p*=0.8175 for Slc6a17^+/+^ vs. Slc6a17^P633R/P633R^, *p*=0.9804 for Slc6a17^P633R/+^ vs. Slc6a17^P633R/P633R^); Pro (D, *p*=0.9674 for Slc6a17^+/+^ vs. Slc6a17^P633R/P633R^, *p*=0.9989 for Slc6a17^P633R/+^ vs. Slc6a17^P633R/P633R^); Leu (E, *p*=0.7116 for Slc6a17^+/+^ vs. Slc6a17^P633R/P633R^, *p*=0.6696 for Slc6a17^P633R/+^ vs. Slc6a17^P633R/P633R^); Gly (F, *p*=0.7376 for Slc6a17^+/+^ vs. Slc6a17^P633R/P633R^, *p*=0.9971 for Slc6a17^P633R/+^ vs. Slc6a17^P633R/P633R^); Ala (G, *p*=0.9871 for Slc6a17^+/+^ vs. Slc6a17^P633R/P633R^, *p*=0.986 for Slc6a17^P633R/+^ vs. Slc6a17^P633R/P633R^); Gln (H, *p*=0.0029 for Slc6a17^+/+^ vs. Slc6a17^P633R/P633R^, *p*=0.0117 for Slc6a17^P633R/+^ vs. Slc6a17^P633R/P633R^); Cys (I, *p*=0.8143 for Slc6a17^+/+^ vs. Slc6a17^P633R/P633R^, *p*=0.5475 for Slc6a17^P633R/+^ vs. Slc6a17^P633R/P633R^); Met (J, *p*=0.9942 for Slc6a17^+/+^ vs. Slc6a17^P633R/P633R^, *p*=0.995 for Slc6a17^P633R/+^ vs. Slc6a17^P633R/P633R^); Ser (K, *p*= 0.9955 for Slc6a17^+/+^ vs. Slc6a17^P633R/P633R^, *p*=0.8355 for Slc6a17^P633R/+^ vs. Slc6a17^P633R/P633R^); His (L, *p*=0.6177 for Slc6a17^+/+^ vs. Slc6a17^P633R/P633R^, *p*=0.4316 for Slc6a17^P633R/+^ vs. Slc6a17^P633R/P633R^).

LC-MS analysis of contents from SVs showed a significant decrease of Gln level in Slc6a17^P633R/P633R^, as compared to Slc6a17^+/+^ and Slc6a17^P633R/+^ mice (Figure 10H). The difference in Gln between Slc6a17^+/+^ and Slc6a17^P633R/+^ was not statistically significant (Figure 10H). The levels of Glu (Figure 10A), GABA (Figure 10B) and ACh (Figure 10C) in SVs were not significantly different among Slc6a17^+/+^, Slc6a17^P633R/+^ and Slc6a17^P633R/P633R^ mice. The levels of the other 8 AAs in the SVs were also not significantly different among Slc6a17^+/+^, Slc6a17^P633R/+^ and Slc6a17^P633R/P633R^ mice (Figure 10D, 10E, 10F, 10G, 10I, 10J, 10K and 10L).

Thus, mislocalization of SLC6A17^P633R^ outside the SVs correlates with the inability of Gln enrichment in SVs (Figure 10H), as well as behavioral impairment (Figure 3 D, F, H and I), establishing the significance of SLC6A17 localization on the SVs.

## Discussion

We have taken multiple approaches to analyze 7 kinds of genetically modified mice (4 modified in the germline and 3 in adult brains) and our results indicating that Gln as an endogenous substrate of SLC6A17 and SLC6A17 is necessary and sufficient for Gln presence in SVs in vivo.

We have discovered the presence of Gln in the SVs and shown that it is functionally important because two mutations pathogenic for human ID patients both resulted in reduction of Gln in the SVs. We discuss these findings in basic neurobiology with implications in human ID pathology.

### The discovery of Gln in SVs and its functional significance

With behavioral studies in two types of Slc6a17 mutant mice, we have shown that one ID pathogenic Slc6a17 mutation (Slc6a17^P633R^) causes defective learning and memory (Figure 3), similar to the phenotype of Slc6a17 knockout mice (Figure 2). With biochemical purifications and genetically assisted EM, we have confirmed SLC6A17 localization on SVs (Figure 4).

We demonstrate the functional significance of the SV localization of SLC6A17 by showing that the human ID mutation Slc6a17^P633R^ causes mislocalization of SLC6A17 outside the SVs (Figure 3B and S6A). Gln transport into the SVs was decreased in Slc6a17^P633R^ mutant mice (Figure 10I).

With multiple rounds of biochemical purification of SVs and chemical analysis of SV contents of genetically modified mice, we have found that Gln is present in SVs containing SLC6A17 and that SLC6A17 is sufficient for Gln presence in SVs (Figure 5, 6 and 7). With two lines of germline mutant mice and virally mediated gene edited mice, we demonstrate that Gln is the only molecule reliably and reproducibly reduced in SVs when Slc6a17 gene is deleted (Figure 8 and 9).

We have also characterized mice carrying a human ID mutation SLC6A17^G162R^ and found it could not transport Gln into SVs (Figure 7), further supporting the importance of Gln in SVs. Thus, two mutations pathogenic in human ID are both LOF mutations: SLC6A17^P633R^ with mislocalization outside the SVs and the SLC6A17^G162R^ with defective Gln transport into the SVs.

The fact that all four kinds of SLC6A17 LOF mutations (two KOs, two point mutations) have led to reduction of Gln in SVs strongly support the functional significance of Gln in SVs.

### Functional mechanisms of Gln in SVs

There are three possible functions for Gln in SVs: serving as a neurotransmitter, serving as a carbon source and serving in a function so far unknown.

The first possibility is most attractive. SLC6A17 is a member of the SLC6 family, which is also known as the NTT family because many members of this family are transporters for neurotransmitters (Blakely and Edwards, 2012; Broer, 2006; Kristensen et al., 2011; Rudnick et al., 2014): SLC6A1 for GABA (Guastella et al., 1992; Clark et al., 1992; Borden et al., 1992; Lopez-Corcuera et al. 1992), SLC6A2 for noradrenaline (Pachoczyk, Blakely and Amara, 1991), SLC6A3 for DA (Giros et al., 1991; Kilty et al., 1991; Shimada et al., 1991), SLC6A4 for 5-HT or serotonin (Blakely et al., 1991; Hoffman et al., 1991), SLC6A5 and SLC6A9 for Gly (Guastella et al., 1992; Smith et al. 1992; Liu et al., 1992). Most of these are cytoplasmic transporters for transporting neurotransmitters from the synaptic cleft into the presynaptic cytoplasm.

The presence of the SLC6A17 protein on the vesicular membrane further supports the possibility that it is a neurotransmitter transporter, because most of the vesicular transporters with known substrates transport neurotransmitters. If this is true for SLC6A17, then Gln would be a good candidate for a new neurotransmitter. However, this exciting possibility remains to be investigated. For Gln to be a transmitter, in addition to its presence in the SVs, more criteria should be satisfied: 1) it should be released from the nerve terminals upon electric stimulation, 2) it should be active on postsynaptic membrane, 3) it should be up-taken by a cytoplasmic transporter back into the presynaptic terminal or be removed otherwise (such as degraded enzymatically), and 4) a pharmacological blocker inhibiting its activity should interfere with chemical transmission in a physiologically significant context. Because Gln is abundant both intracellularly and extracellularly, we expect that demonstration of all the above would take some time, and likely by many laboratories.

There is a Glu/Gln cycle in which Glu released from neurons is taken up by glia and transformed into Gln, followed by Gln release from glia and taken up by neurons and transformed into Glu (Bak et al., 2006; Benjamin and Quastel, 1972; Ottersen et al., 1992; van den Berg and Garfinkel, 1971). Gln uptake across the neuronal cytoplasmic membrane relies on system A transporters (Mackenzie et al., 2003; Reimer et al., 2000; Varoqui et al., 2000; Weiss et al., 2003), while Gln uptake across the glia cytoplasmic membrane uses system L (Deitmer et al., 2003; Kanai et al., 1998), system N transporters (Broer and Brookes, 2001; Chaudhry et al., 2002; Chaudhry et al., 1999) and system ASC (Broer et al., 2000). It is unlikely that SLC6A17 and the Gln it transports could be part of the Glu/Gln cycle, because: 1) the Glu/Gln cycle uses cytoplasmic transporters for Gln, not vesicular transporters for Gln; 1. 2) Gln is transformed into Glu in the cytoplasm of neurons before being transported in the SVs as Glu; 3) in all of our experiments presented here, when SLC6A17 was genetically disrupted, Gln was consistently decreased, but Glu was either unchanged but not concomitantly increased, which was against the expectation of the Glu/Gln cycle. The only result in which Glu was increased when SLC6A17 was decreased came from an experiment with virally mediated genetic knockdown of SLC6A17 (Figure 9E), but GABA was also increased in that experiment (Figure 9F). When the same experiment was carried with a germline genetic knockout of SLC6A17, Glu was not changed when SLC6A17 was decreased (Figure 9A).

The possibility of Gln as a carbon source is based on the fact that it is one of the major carbon sources (Hensley et al., 2013; Reitzer et al., 1979). It would be very curious if a carbon source should be stored in the SVs. It would suggest an entirely new function for Gln in the nerve terminal which has never been suspected before and may require considerable efforts before one can fully understand.

Multiple other functions for Gln have been proposed (i.e., Curi et al., 2005; Wischmeyer, 2006).

### Other substrates for SLC6A17

Of the nine AAs (Ala, Gly, Leu, Pro, Cys, Gln, Gly, His and Ser) previously reported to be in vitro substrates for SLC6A17 transport in cell lines (Parra *et al*., 2008; Zaia and Reimer, 2009), we have reliably and reproducibly found evidence supporting only one (Gln) as present *in vivo* in all 5 types of experiments conducted with SV purifications: SVs immunoisolated from the WT brains by anti-Syp (Figure 5), SVs immunoisolated from the brains of mice with the HA tag fused in frame to the endogenous SLC6A17 protein (Figure 6), SVs immunoisolated from mice with virally introduced overexpression of Syp-HA, SVs immunoisolated from mice with virally introduced overexpression of SLC6A17^G162R^-HA, and SVs immunoisolated from mice with virally introduced overexpression of SLC6A17-HA (Figure 7). While we did observe a moderate increase of His in one set of experiments (OE SLC6A17-HA vs WT), we did not find evidence for its changes in other experiments.

Because multiple transporters can transport AAs, and the relative contribution of each transporter varies and the relative abundance of each small molecules and their detectability varies, we can not rule out the possibility that SLC6A17 can contribute to the transport of other AAs into the SVs. However, our results can establish that Gln is transported *in vivo*.

In summary, integration of genetics, biochemistry, behavioral analysis, mass spectrometry and electron microscopy has resulted in the discovery of Gln as a novel chemical in SVs, provided *in vivo* evidence that SLC6A17 is necessary and sufficient for Gln in SVs, generated animal models for, and suggested pathogenetic mechanism of, human ID. However, while we have answered the question how Slc6a17 mutations may cause ID in animals, our research has also raised more questions. This natural process of questions begetting answers begetting questions is intrinsic to scientific inquiries and is a source of endless fun in conducting research.

## Materials and methods

### Reagents

Chemical reagents for biochemical purification and immunoisolation were all ordered form Sigma-Aldrich. The following antibodies were purchased from Synaptic System mouse anti-Synaptophysin 1 (cat.101011 for IP), rabbit anti-Synaptophysin 1 (cat.101002 for IB), rabbit anti-Synaptotagmin 1 (cat.105008 for IB), rabbit anti-VGLUT 1 (cat.135302 for IB), rabbit anti-VGAT (cat.131002 for IB), rabbit anti-VGLUT 2 (cat.135402 for IB), rabbit anti-Proton ATPase (cat.109002 for IB), rabbit anti-Synaptobrevin 2 (cat.104202 for IB), rabbit anti-SV2A (cat.119002 for IB), rabbit anti-SNAP23 (cat.111002 for IB), mouse anti-GluN1(cat.114011 for IB), and rabbit anti-ERC 1b/2 (cat.143003 for IB). The following antibodies were purchased from Abcam: mouse anti-Synaptophysin (cat. ab52636 for IF), mouse Alexa Fluor® 488 anti-Synaptophysin (cat. ab196379 for IF), mouse anti-PSD95 (cat.ab76115 for IB), rabbit anti-Cathepsin D (cat.ab75852 for IB), mouse anti-PSMC6 (cat.ab22639 for IB), rabbit anti-GLUT4 (cat.ab33780 for IB), and rabbit anti-Transferring receptor (cat.ab84036 for IB). The following antibodies were purchased from Cell Signaling Technology: rabbit anti-LAMP2 (cat.49067 for IB), rabbit anti-EEA1 (cat.3288 for IB), rabbit anti-ERp72 (cat.5033 for IB), rabbit anti-Goglin-97 (cat.1292S for IB), rabbit anti-GM130 (cat.12480 for IB), rabbit anti-VDAC (cat.4661S for IB), rabbit anti-Syntaxin 6 (cat.2869 for IB), rabbit anti-LC3B (cat.2775 for IB), and rabbit anti-HA (cat.3724S for IF or IB). Mouse anti-FLAG M2 HRP conjugated antibodies (cat.A8592 for IB) were purchase from Sigma.

### Animals

All animal procedures were approved by the Animal Center of Peking University, and the experiments were carried out in accordance with the guidelines of Institutional Animal Care and Use Committee (IACUC) of Peking University. Mice were weaned at the age of 21 days. Mice were maintained under standard conditions (12 h light, 12 [dark schedule). Room temperature was 23[[C. Humidity was 40–60%. Mice for behavior assays were 10-16 weeks old.

All transgenic mouse lines were generated in C57BL/6J background. Ai14 mice were a gift from Dr. Hongkui Zeng. Rosa26-LSL-Cas9 mice were obtained from The Jackson Laboratory (028551). SLC6A17-2A-CreERT2 and SLC6A17-KO mice were generated by CasGene biotech (Beijing, China). SLC6A17-HA-2A-iCre and SLC6A17P633R mice were generated by Biocytogen (Beijing, China). Mutant strains were generated by CRISPR/Cas9. Approximately 1.5 kb homologous arms upstream and downstream of target site were cloned into the targeting vector. The cleavage efficiency of single guide RNAs (sgRNAs) targeting SLC6A17 were tested in HEK cells. The sgRNAs selected for final injection were: 5’- ccaacggacgctatggaagcggc-3’ for SLC6A17-2A-CreERT2 mice; 5’-catgcccaggtaacacctatggg-3’ and 5’-cctgttaacaacactatctgaca-3’ for SLC6A17-KO mice; 5’-actgaggggtgctggccaagagg-3’ for SLC6A17-HA-2A-iCre mice; and 5’-ctgctagaatccaggaggcagg-3’ for SLC6A17P633R mice. All mutant strain were confirmed by PCR sequencing of the targeting regions. Southern blots were performed by Biocytogen for SLC6A17-HA-2A-iCre and SLC6A17P633R mice to rule out random insertions. For CreERT2 induction, tamoxifen was administrated intraperitoneally (i.p.) 1mg/day/mice for 5 consecutive days at the age of 4-5 weeks. Two weeks after tamoxifen injection, mice were killed for histology examination.

### Histology and Immunocytochemitry

Mice were anesthetized, perfused, and post-fixed as described previously (Liu, et al., 2011; Qin and Luo, 2009). Briefly, mice were deeply anesthetized with Tribromoethanol (250 mg/kg; i.p.) and transcardially perfused with cold 0.1M PBS followed by 4% paraformaldehyde (PFA) in 0.1 M PBS. Brains were fixed overnight in 4% PFA at 4 °C and then immersed in 30% sucrose until it sank to the bottom at 4 °C. It was embedded in the Tissue-Tek O.C.T. compound and frozen in a freezing microtome chamber (Leica CM 3050; Leica). Sections were 40 μm thick and collected on adhesion glass slides (P/N.80312-3161, CITOTEST). Slc6a172A-CreERT2::Ai14 mice sections were washed with 0.3% Trtion X-100 in PBS (PBST) and then placed under cover-slips with a mounting solution (Fluoroshield with DAPI, F6057, sigma). Sections were imaged by confocal microscope (Zesis LSM710) or Slide scanner (Zeiss Axio Scan. Z1). For immunofluorescence, freshly prepared slides were washed with PBST, followed by 1 h 10% normal goat serum (in PBST) to block non-specific binding. Primary antibodies were diluted to working concentrations in PBST, and incubated with slides overnight at 4°C. Slides were washed with PBST and incubated with secondary antibodies for 1h at room temperature. After PBST washes, slides were placed under cover-slips with a mounting solution. For double staining in Figure 3B, S7B and S7C, anti-HA antibody (RmAb, 1:200 dilution) was applied and labeled in Alexa Fluor 546, then Alexa Fluor 488 Anti-Syp (RmAb, ab196379, Abcam, 1:200 dilution) was applied. Slides were placed under cover-slips with a mounting solution and images were analyzed by confocal microscope (Zesis LSM710).

### Behavioral Tests

Mice were placed in the experiment room for at least 1 h before behavioral tests. All experiments were performed with the genotypes blind to the researchers. 70% ethanol was used to clear odor cues after each test.

#### Open field Tests

Experiments were conducted in a square Plexiglas apparatus (40× 40× 35 cm) with illumination kept at 100-200 lux. Mice were gently placed in the center of the apparatus and allowed to explore for 30 minutes (min). Activities were recorded by a digital camera (HDR-XR550, SONY) set directly above the apparatus. After each trial, the apparatus was cleaned and the animals put back to home cage. Videos were analyzed by a home-made program created for SLC6A17-KO mice and Open Filed (Mobile Datum, Shanghai, China) for SLC6A17P633R mice. The home-made program is based on OpenCV. Briefly, mice were extracted by subtracting background which was updated for each frame in order to prevent environmental light shift. The center of the mouse was calculated, from which the distance was determined.

#### Novel object recognition task

The experimental protocol was adopted from previous report (Bevins and Besheer, 2006). On the first day, mice were individually habituated in an open box (40× 40× 25 cm). Next day, two identical objects (O) were placed into left and right corners of the box and mice were allowed to explore for 10 min. Object recognition was tested 1 hr later. One familiar object (O) was replaced by a novel object (N) before mice were placed back into the box. Mice were allowed to explore freely for 5 mins. The time spent in the exploring the object was measured. Discrimination ratio was calculated by novel object interaction time/total interaction time with both objects. Testing sessions were recorded by a digital camera (HDR-XR550, SONY) and analyzed by software (Mobile Datum, Shanghai, China).

#### Morris water maze

The Morris water maze task was modified according to previously described procedures (Hitti and Siegelbaum, 2014; Qin et al., 2017). The maze contained a round pool (diameter: 120 cm) and a round platform (diameter: 10 cm, 2 cm below the water surface). Water was 50 cm deep and set at 20 ± 1 °C. Visual cues were placed above the pool. Titanium white was added into water to make it opaque for mice. We trained mice for four 1 min-trials per day. If a mouse successfully found the platform, it was taken back to home cage 15 s after locating the platform. If a mouse failed to find the platform during 1 min, it was gently placed on platform for 15s then taken back to the home cage. On days 1 and 2, cued learning was conducted to find the platform marked with a visible object. Mice were trained from different start points at each trial. On days 3 to 7, the object was removed while mice were trained with different start points to locate the hidden platform whose location was different with cue learning session. The latency for each animal to find the platform was recorded. On day 8, spatial memory was tested with a 1-min probe trial in which the platform was removed. Crossings into the platform area and total time spent in the target platform quadrant were recorded. The trajectories of probe trial were analyzed by Noldus Video Tracking Software (Ethovision XT 15, Noldus, Netherlands).

#### Fear conditioning

A three-day delay fear-conditioning protocol was employed to test fear memory (Hitti and Siegelbaum, 2014; Qin et al., 2017). A conditioning chamber (25 cm × 25 cm × 40 cm) with 2 types of contexts (context a: black walls, metal net floor for electrical stimulation; context b: white walls and white plastic floor) was used. On day 1, mice were trained in context a. Two rounds of training were performed. Each round consisted of a 1 min adaptation of environment, followed by a cued tone (30 s, 300 Hz, 90 dB sound) and a foot shock (2 s, 0.75 mA constant current). On day 2, contextual fear memory was assayed by placing the mice back in context a for 300 s. On day 3, mice were placed in context a. After 180 s, a training tone was sounded for 180 s. 5% acetic acid were used to clean chambers between day 3 and day 2. Two kinds of apparatus were used. SLC6A17-KO mice were tested in Fear Conditioning (Mobile Datum, Shanghai, China) and SLC6A17P633R mice were tested in Startle and Fear Combined System (Panlab, Spain). Animal activities were recorded and the percentage of time spent freezing (defined as the absence of all movement except for respiration) was measured by the software accordingly. Fear Conditioning (Mobile Datum) is a video-based recording system, so freezing time was re-analyzed by a double blinded researcher to revise software variations. Startle and Fear Combined System (Panlab) is a gravity-based recording system.

#### Elevated Plus Maze

The experimental protocol was adopted from previous reports (Kouser, et al., 2013; Powell et al., 2004). Mice were placed in the center of a white plastic elevated plus maze, each arm was 33 cm long and 5 cm wide, with 15-cm-high opaque plastic walls on closed arms. Mice were allowed to explore for 5 min, under the white light (∼20 lux). Animal activities were recorded and analyzed by a software (Mobile Datum, Shanghai, China).

### Subcellular Fractionation

Experimental protocols were based on previously described methods for SV purification (Evans, 2015; Huttner et al., 1983). Homogenate buffer [0.32 M sucrose and 5 mM HEPES-NaOH, pH 7.4, protease inhibitor cocktail (Complete EDTA-free, Roche)] was pre-cooled on ice, and all operations were conducted on ice. Whole mouse brains were homogenized (kept as fraction H) in the homogenate buffer in a Potter Elvehjem homogenizer (Wheaton Instruments, US, 9 strokes at 900 rpm) and centrifuged at 1000 g for 10 min. The supernatant was kept as Fraction S1 and pellet containing large membranes was resuspended (kept as Fraction P1). Fraction S1 was then centrifuged at 13,000 g for 20 min to yield a crude synaptosomal pellet (kept as Fraction P2), and remaining supernatant collected as Fraction S2. Synaptosomes in Fraction P2 were osmotic-lysed by adding 9 volumes of ice-cold ddH_2_O (protease inhibitor cocktail added), followed by quick homogenization (3 strokes, 2000 rpm) and 1/20 volume 1M HEPES-NaOH (pH 7.4) was added. The osmotic-lysed samples were incubated on ice for 40 min to maximize yields. Lysates were then centrifuged at 25,000 g for 20 min. Supernatant was collected as Fraction LS1 and pellet containing synaptic plasma membranes was resuspended (kept as Fraction LP1). Fraction LP2 was obtained after centrifugation of fraction LS1 for 2 h at 165,000 g in a SW40 Ti rotor (Beckman, US). Corresponding supernatant was collected as Fraction LS2. Further purification of the SVs in Fraction LP2 was performed by centrifugation on a continuous sucrose gradient. Fraction LP2 was resuspended in gradient buffer (40 mM sucrose, 5 mM HEPES-NaOH, pH 7.4) and 200 μl (1 mg/ml) resuspended fraction LP2 was layered onto a 11 ml 50-800 mM continuous sucrose gradient. The gradient was centrifuged at 65,000 g for 5 h in SW40 Ti rotor and then divided to 11 equal fractions from the top to the bottom. 4 volumes of pre-cooled methanol (-20°C) was added into sub-fractions to precipitate proteins. Proteins were collected by centrifugation in 20,000 g for 20 min. All centrifugations were conducted at 4°C.

### Immunoisolation of SVs

Experimental protocols were adopted from previous established methods (Burger, et al., 1991; Burger et al., 1989; Martineau et al., 2013). The entire experiments were conducted at 2-8 °C. Whole brains were homogenized immediately after decapitation by ice-cold immunoisolation buffer (4 mM HEPES-KOH, 100 mM K2-tartrate, 2 mM MgCl2, pH 7.4, and protease inhibitor cocktail) in Potter Elvehjem homogenizer (15 strokes, 2000 rpm). Homogenates were centrifuged 35,000 g for 25 min at 4 °C. The supernatant (protein concentration 3 mg/ml) was used as input to incubated with magnetic beads coupled with antibody. For anti-Syp immunoisolation, anti-Syp (101 011, SySy) antibody was pre-incubated with Pierce protein G magnetic beads (88848, Thermo Scientific) in 0.1% BSA PBS at 4 °C overnight. The anti-Syp antibody and mice IgG (02-6502, Invitrogen) were bound to Pierce protein G magnetic beads at the ratio of 33 μg of IgG per mg beads. For anti-HA immunoisolation, Pierce anti-HA magnetic beads (88837, Thermo Scientific) were used and washed just before each experiment. Beads were incubated with input supernatant for 2 h with constant rotation at 4°C. Beads were washed 6 times by immunoisolation buffer after incubation. Following the final wash, beads were magnetically separated and vesicular contents were extracted by 50 μl 80%/20% methanol/H_2_O (-80 °C) with 5 μM internal standards. To maximize the extraction efficiency and protein precipitation rate, the extraction mix was transferred to -20 °C for 1 h. The extraction mix was then centrifuged for 20 min at 20,000 g, and supernatant was transferred to new vials subjected to LC-MS. The remaining beads were extracted by SDS loading buffer for immunoblot analysis.

### Immunoblotting

Proteins were extracted from each sample with SDS loading buffer (3% SDS, 15% glycerol, 180 mM Tris-HCl, 4% β-mercaptoethanol). For subcellular fractionation samples, protein concentrations were determined by the BCA protein assay (Pierce, 23250, Thermo Scientific), and equal quantities of proteins were subjected to SDS-PAGE. For immunoisolation samples, equal volumes of SDS loading buffer were added to magnetic isolated beads. Samples were incubated in 75 °C for 20 min to better immunoblot transmembrane proteins. Briefly, samples were loaded to 10% SDS-PAGE gels (TGX FastCast Acrylamide Kits, 1610183, Bio-Rad) and transferred to 0.45mm NC membranes (HATF00010, Millipore) for 2 h. Membranes were subjected to 5% milk in TBST buffer (Tri-buffered Saline with Tween 20) at room temperature for 1h to block non-specific binding. Primary antibodies were prepared in TBST and incubated with membranes at 4 °C overnight. Secondary antibodies were either anti-rabbit IgG HRP (A6154, Sigma, 1:5000 dilution) or anti-mouse IgG HRP (715-035-150, Jackson Immuno Research, 1:5000 dilution). Final results were visualized using chemiluminescence (Merck Millipore) in e-BLOT Touch Imager (e-BLOT Life Science, Shanghai).

### AAV Plasmid Construction and Virus Injection

AAV-PHP.eb viruses were packaged in HEK293T cells by the vector center of CIBR. The titres of AAVs were determined by real-time PCR. Viruses were subpackaged and stored in -80 °C before use. All AAV constructs used for viral packaging were linearized by restriction enzymes to insert target sequences. Bacterial cultures were grown at 33 °C. AAV-PHP.eb viruses were packaged by Vector Center in CIBR. Viruses were delivered retro-orbitally into the venous sinus of 6-week-old mice. For APEX2 staining, 2*1011 particles of virus were injected. For gene overexpression and sgRNA expression, 1*1011 particles of virus were injected.

**Construction of pAAV-hSyn-SLC6A17-APEX2-WPRE-pA plasmid:** this plasmid consisted of human SLC6A17 cDNA, 3×HA tag, V5 tag, and APEX2. SLC6A17 cDNA was cloned from commercially available cDNA library (Vigene, Shandong, China). The 3×HA tag (YPYDVPDYAGYPYDVPDYAGSYPYDVPDYA) were chosen the same version as SLC6A17-HA-iCre mice was in-frame fused with cDNA by GS4 linker (GGGGSGGGGS GGGGSGGGGS). V5-APEX2 sequence were cloned from mito-V5-APEX2 plasmid (#72480, addgene). V5-APRX2 were first cloned into pAAV-hSyn-WPRE-pA plasmid. Then SLC6A17-HA was inserted before V5-APEX2.

**Construction of pAAV-hSyn-SLC6A17-HA-WPRE-pA plasmid and pAAV-PHP.eb:hSyn-SLC6A17G162R-HA-WPRE-pA plasmid:** SLC6A17-HA sequence consists of pAAV-hSyn-SLC6A17-APEX2. G162R point mutation was introduced by PCR (forward primer: 5’-attataatgtgatcatcAggtggagcatcttc-3’; reverse primer: 5’-Tgatgatcacattataatacagcccc-3’) in a middle plasmid without ITR element. Then the SLC6A17G163R-HA sequence was cloned into pAAV-hSyn-WPRE-pA plasmid.

**Construction of pAAV-hSyn-DIO-Syp-HA-WPRE-pA-U6-3-t::gRNA WPRE-pA plasmid:** tRNA-sgRNA design was modified from 1,2. sgRNAs were chosen based on published database3. The sequence of selected gRNA: sgRNA1 (5’- CGATGCTCCAGGCCACAAGGAGG -3’, targeting exon 5); sgRNA2 (5’-GCCGTGGCAGCATTGGTGTGTGG -3’, targeting exon 3); sgRNA3 (5’-TGGGCCTGGGCAACATCTGGAGG -3’, targeting exon 2). Syp-HA sequence was synthesized and cloned into AAV backbone with hSyn promoter and DIO cassette.

### Electron Microscopy with APEX2

Procedures for APEX2 based EM labeling were modified from previous reports (Lam, et al., 2015; Martell et al., 2017). Briefly, the AAV injected animals were anesthetized and perfused with PBS and 4% paraformaldehyde (PFA) and 1% glutaraldehyde (GA) for pre-fixation. Mouse brains were dissected and cut into approximately 1 mm3 blocks and was added into 2.5% GA for post-fixation for 2 h. Samples were then washed with 0.1 M phosphate buffer (PB). Following pre-incubation in 0.5 mg/ml 3,3-diaminobenzidine (DAB, D5905, Sigma) for 30 mins, 0.03% H_2_O_2_ (final concentration) was added into samples for APEX2 oxidation (Note that different sources of DAB had considerable disparity, and D5905 produced the best signal than other DABs in our lab), although the recommended D8001 was poorer than D5905) (Ludwig, 2020; Martell et al., 2017). The reaction was monitored by the dark brown color in tissue block and was stopped by removing the DAB solution and washed three times with 0.1 M PB. Most of the successful DAB staining usually turned deep color within 10 mins. After DAB staining, samples were incubated in 4% osmium tetroxide in 0.1M PB on ice for 1 h in dark environment. Samples were washed with ddH_2_O and incubated in 1 % aqueous uranyl acetate overnight at 4°C. Next day, samples were dehydrated in graded ethanol series (30%, 50%, 75%, 85%, 95% and 100%) at 4°C and rinsed once in acetone with each rinse for 10 mins at room temperature. Samples were infiltrated in Epon 812 resin using a 1:3, 1:1 and 3:1 ratio of resin and acetone for 2 h each followed by 3 infiltrations with 100% resin for 30 mins each. Finally, samples were embedded in fresh resin in a vacuum oven for 24 h at 65°C. Blocks were sectioned by UC7 ultramicrotome (Leica Microsystems, Vienna, Austria) into 70 nm. A 120 kV JEOL JEM-1400Flash TEM (JEOL, Japan) was used to image the brain and images were acquired by an XAROSA camera (Emsis GmbH, Muenster, Germany). Images were saved in a .tif format and were further analyzed using FIJI-ImageJ 2.1.0/1.53c (Wayne Rasband, NIH). The segmentation of SVs was performed using the TrakEM2 v1.0a (Univ/ETH Zurich) plugin in FIJI and the mean intensity of DAB + or - SV was quantified using a custom-made macro. Figure were created using OriginPro 2022.

### LC-MS for Chemical Detection

LC-MS was used to detect and quantify the chemical contents of samples based on previous reports (Abu-Remaileh, et al., 2017; Contrepois et al., 2015; Mackay et al., 2015; Wernisch and Pennathur, 2016; Zhang et al., 2014; Zhang et al., 2012). A Vanquish UHPLC system coupled to a Q Exactive HF-X mass spectrometer (both instrument Thermo Fisher Scientific, USA) were used for LC-MS analysis along with SeQuant ZIC-HILIC column (150 mm×2.1 mm, 3.5 μm, Merck Millipore, 150442) in the positive mode and SeQuant ZIC-pHILIC column (150 mm×2.1 mm, 5 μm, Merck Millipore, 150460) in the negative mode. For ZIC-HILIC column, the mobile phase A was 0.1% formic acid in water and the mobile phase B was 0.1% formic acid in acetonitrile. The linear gradient was as following: 0 min, 80% B; 6 min, 50% B; 13 min, 50% B; 14 min, 20% B; 18 min, 20% B; 18.5 min, 80% B; 30 min, 80% B. The flow rate used was 300 μl/min and the column temperature was maintained at 30 °C. For ZIC-pHILIC column, the mobile phase A is 20 mM ammonium carbonate in water, adjusted to pH 9.0 with 0.1% ammonium hydroxide solution (25%) and the mobile phase B is 100% acetonitrile. The linear gradient was as following: 0 min, 80% B; 2 min, 80% B; 19 min, 20% B; 20 min, 80% B; 30 min, 80% B. The flow rate used is 150 μl/min and the column temperature is 25 °C. Samples were maintained at 4 °C in Vanquish autosampler. 3 µL of extracted metabolites were injected for each run. IP samples were subjected to ZIC-HILIC column in positive mode for major metabolites detection, and then subject to ZIC-pHILIC column in negative mode for orthogonal detection.

Calibration was performed prior to analysis using the Pierce^TM^ Positive and Negative Ion Calibration Solutions (Thermo Fisher Scientific) every 7 days. Acetonitrile was LC-MS grade (A955, Thermo Fisher Scientific) and all other regents were LC-MS Optima grade from sigma. Isotope-labeled amino acids (sigma) as internal extraction standards were supplemented into extraction solution. Orbitrap mass spectrometer was operated with the following parameters: scanning in Full MS mode (2 μ cans) from 70 to 1050 m/z at 70,000 resolution, with 4 kV spray voltage (3.5 kV for negative mode), 50 shealth gas (35 for negative mode), 12 auxiliary gas (8 for negative mode), capillary temperature 300C, S-lens RF level 55, AGC target 1E6, and maximum injection time 200 ms.

### TMT-labeled SV Proteomics

Procedures for quantitative SV proteomics were modified from previous reports (Boyken, et al., 2013; Gronborg et al., 2010). Proteins were labeled using TMT10plex Isobaric Label Reagent Set (90113, Thermo Scientific). In short, LP2 fractions were purified form both SLC6A17-KO +/+ and +/- mice, with the final step centrifuged in 230,000 g instead of 165, 000 g. Purified LP2 were stored in -80°C for long-term storage. SVs were further purified by immunoisolation with the anti-Syp antibody coupled to magnetic protein G beads incubated with resuspended LP2 fraction for 2h at 4 °C (LP2 should be repeatedly passed through 28 gauge needles to be fully resuspended). After 6 washes in PBS, SVs were magnetically separated and dissociated form beads by adding lysis buffer [10% SDS in 100 mM TEAB (triethyl ammonium bicarbonate)] to samples. Beads were heated in 70°C for 10 min to fully dissolve proteins and then centrifuged 10 min at 16,000 g to remove beads. Protein concentrations were determined by BCA kits (23227, Thermo Scientific) to ensure the total proteins between 50-100 µg. Equal amounts of proteins were transferred into new tubes and adjusted to 100µL with 100mM TEAB. 5µL 200 mM TCEP (3,3’,3’’-phosphinetriyltripropanoic acid) was added and incubated in 55°C for 1 h. Then 5 µL freshly prepare 375 mM iodoacetamide were added to the samples and incubated for 30 minutes in dark environment at room temperature. Following addition of six volumes (∼600µL) of pre-chilled (-20°C) acetone, proteins were precipitated overnight at -20°C. Next day, samples were centrifuged at 8000 g for 10 minutes at 4°C to precipitate proteins. After acetone removal, 100 µL of 100 mM TEAB were added to resuspend proteins. Trypsin was added at 2.5 µg per sample and incubated at 37°C. Then freshly made TMT Label Reagents (0.8 mg/41µL anhydrous acetonitrile) were added to samples and incubated for 1 h at room temperature. 8 µL 5% hydroxylamine were added to the samples and incubated for 15 minutes to quench the reaction. Equal amounts of each sample were combined in a new centrifuge tube and labeled peptides dried in SpeedVac. Pierce™ High pH Reversed-Phase Peptide Fractionation Kit (84868, Thermo Scientific) were used to clean up and fractionate TMT-labeled peptides before LC-MS/MS. Peptides were identified and analyzed by the Proteomic Center of PKU.

### Statistical Analyses

For statistical comparisons of two groups, we used the Nonparametric Mann-Whitney test to compare two columns of data. For comparisons of more than two groups, we used the One-way ANOVA Brown-Forsythe ANOVA test, and Dunnett’s T3 multiple comparisons test. We used Two-way ANOVA with Tukey’s multiple comparisons test for multiple factor comparation. An α level of 0.05 (two-tailed) was set for significance. All statistical analysis was done using Prism 8 (GraphPad Software).

## Data availability

All data are contained in the article.

## SUPPLEMENTAL INFORMATION

Supplemental information includes 10 Figure.

## ACKNOWLEDGEMENTS

We are grateful to CIMR, CIBR, Peking-Tsinghua Center for Life Sciences, Changping Laboratory, Chinese Academy of Medical Sciences (2019RU003), Shenzhen Bay Laboratory for support.

## Author contributions

YR conceived the idea of searching for new neurotransmitters and supervised the project. XJ JZ and XB tried different approaches to find new neurotransmitters. XJ carried out experiments and supervised technicians to carry out the experiments for this paper. XJ, JZ, XB and SY reproducibly found Gln in SVs in independent experiments. XJ, SL, SY, LJ, WL and RM carried out experiments. XJ and YR wrote the manuscript.

## LEGENDS FOR SUPPLENTAL FIGURE

**Figure S1.**
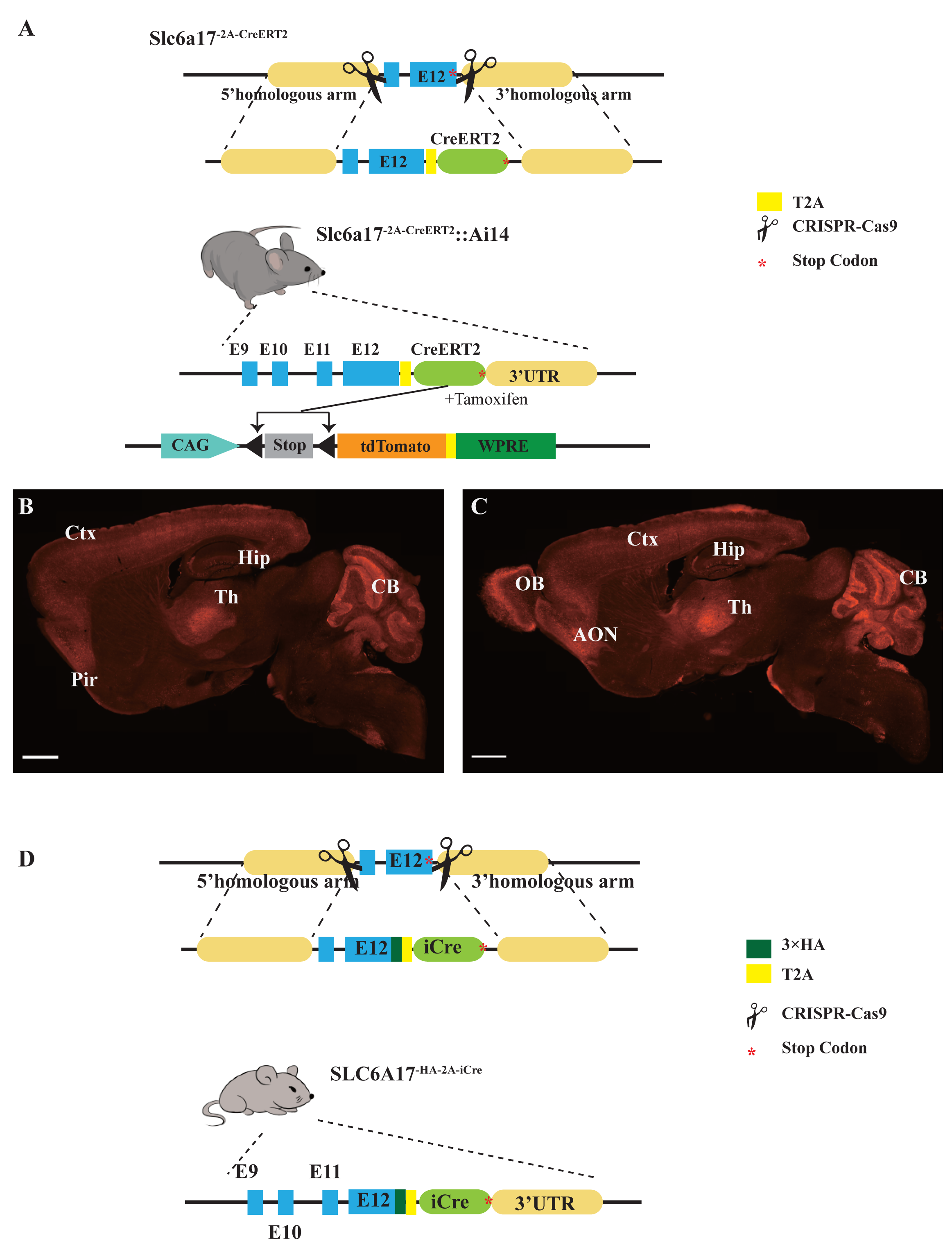
Generation of Slc6a17 Knock-in Mice. (A) A schematic diagram illustrating the knock-in strategy for generating Slc6a17^-2A-CreERT2^ KI mice. This CRISPR/Cas9 mediated strategy allowed CreERT2 to be co-expressed with SLC6A17 protein fused in frame with a T2A sequence. Crossing Slc6a17^-2A-CreERT2^ mice with Ai14 (LSL-tdTomato) mice specifically labeled Slc6a17-expressing cells after tamoxifen injection. (B-C) Representative sagittal sections of Slc6a17^-2A-CreERT2^::Ai14 mouse brains at postnatal day 56. Scale bars=1 mm. (D) A schematic diagram illustrating the knock-in strategy for generating Slc6a17^-HA-2A-iCre^ KI mice. Slc6a17 was fused in-frame with 3 repeats of the HA tag, T2A and iCre.

**Figure S2.**
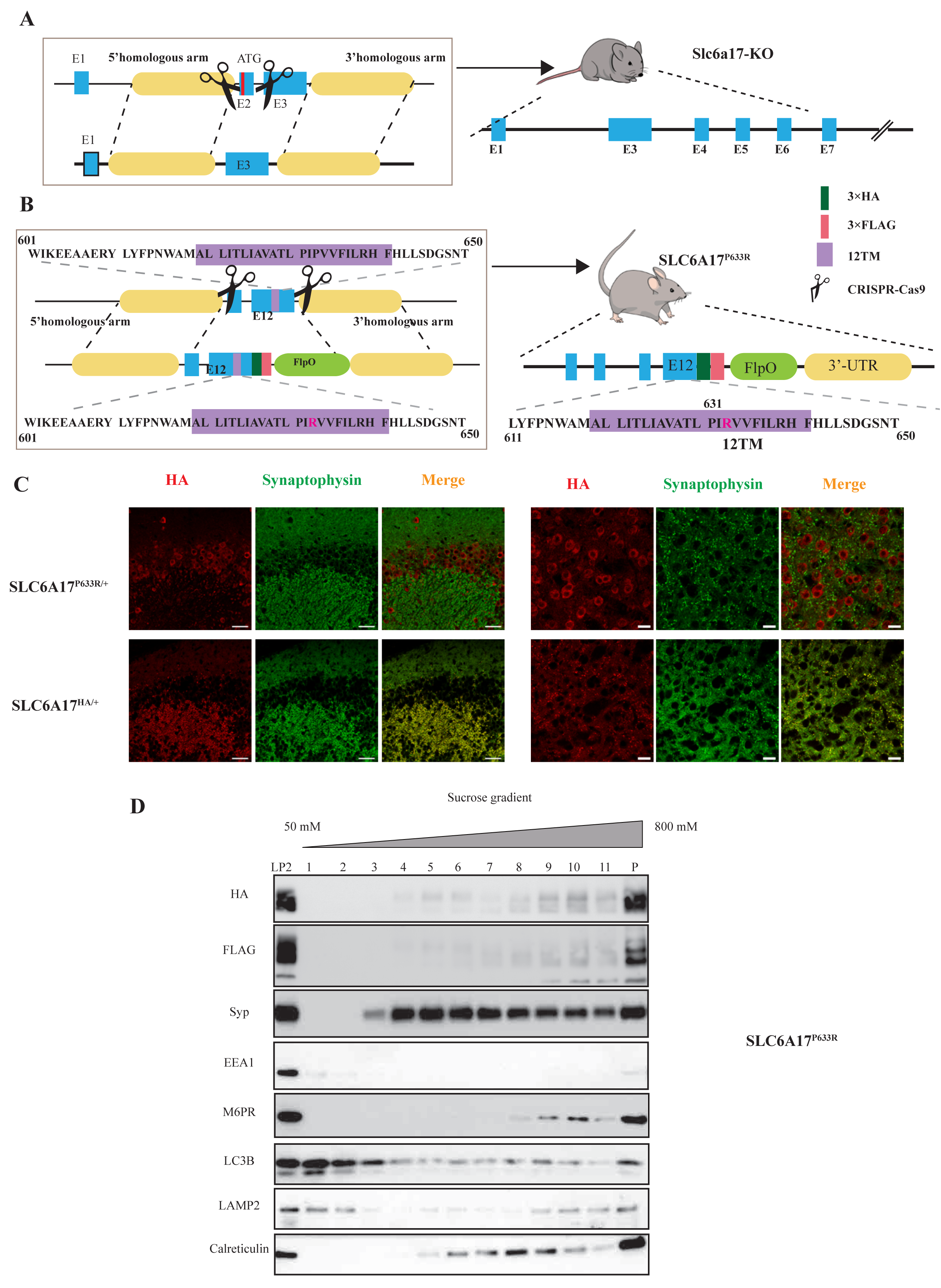
Generation of Slc6a17 Mutant Mice. (A) A schematic diagram illustrating the strategy for the generation of Slc6a17-KO mice by CRISPR/Cas9. Exon 2 encoding the first 99 amino acids of Slc6a17 was deleted. (B) A schematic diagram illustrating the knock-in strategy of generating Slc6a17^P633R^ point mutation mice by CRISPR/Cas9. The mutated exon 12 was exchanged with endogenous exon 12, while both the HA-and the FLAG-tags were fused in-frame to the C terminus of the mutated Slc6a17. (C) Confocal images of the hippocampal CA3 region (left) and anterior thalamus (right) in both Slc6a17^HA/+^ and Slc6a17^P633R/+^ mice after immunocytochemistry with anti-Syp and anti-HA antibodies. Scale bar=10 μm. (D) Sucrose gradient analysis of Slc6a17^P633R^-HA LP2 fraction. SLC6A17^P633R^ protein is indicated by HA and FLAG tags. HA and FLAG signals were mostly present in the LP2 input but not in layers 2-5 of the sucrose gradient, which were SV rich as shown by Syp and Syb2 positive immunoreactivities. On the other hand, SLC6A17^G162R^ protein is still present in layers 2-5.

**Figure S3.**
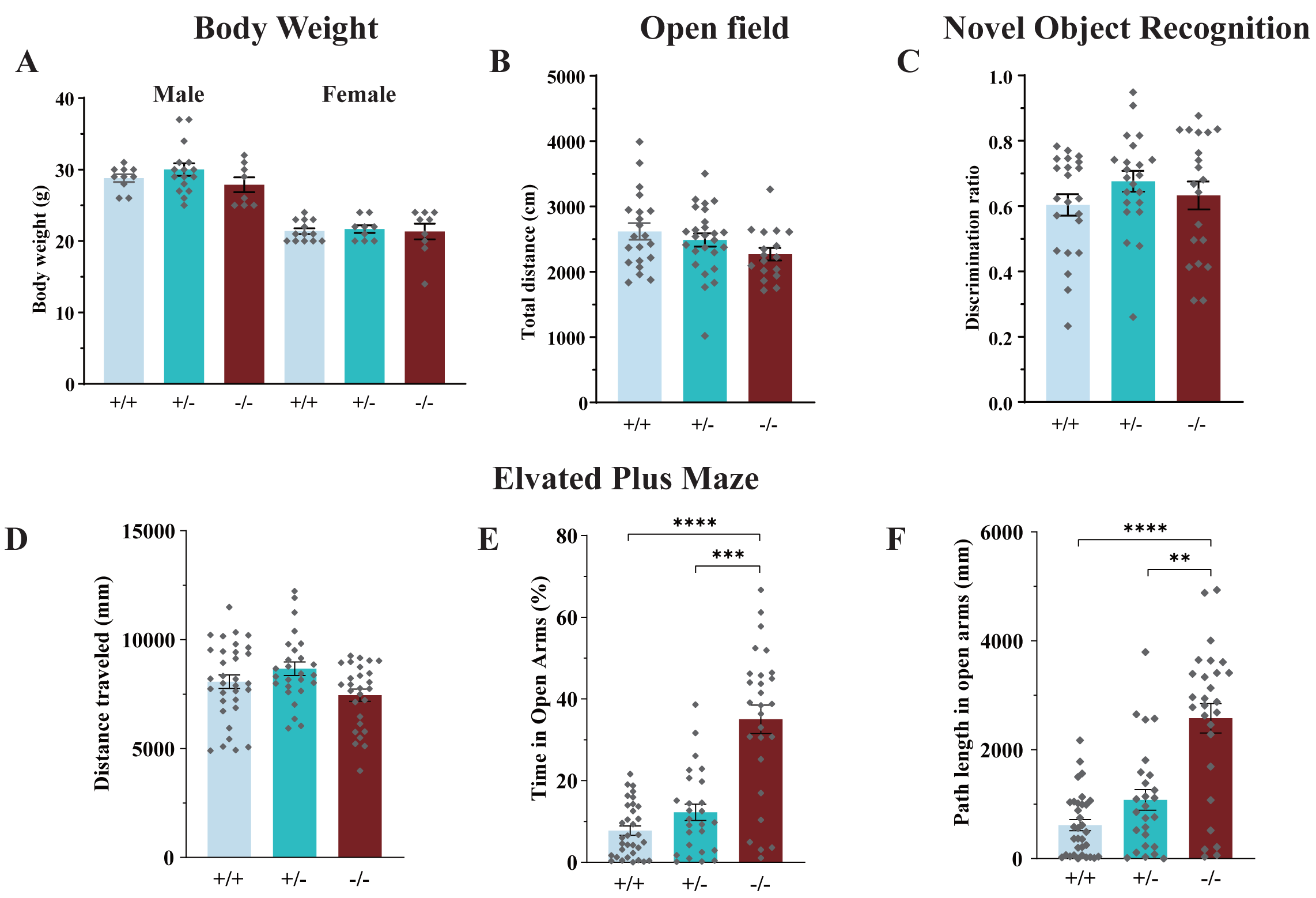
Phenotypical Analysis of Slc6a17-KO Mice. (A) Slc6a17-KO mice were not significantly different from the WT in body weight at the age of 10 week, regardless of sex (male: n=10, 16, 8 for Slc6a17^+/+^, Slc6a17^+/-^, Slc6a17^-/-^, respectively, *p*=0.8147 for Slc6a17^+/+^ vs. Slc6a17^-/-^, *p*=0.3456 for Slc6a17^+/-^ vs. Slc6a17^-/-^; female: n=13, 9, 9 for Slc6a17^+/+^, Slc6a17^+/-^, Slc6a17^-/-^, respectively, *p*> 0.9999 for Slc6a17^+/+^ vs. Slc6a17^-/-^, *p*=0.9896 for Slc6a17^+/-^ vs. Slc6a17^-/-^). (B) Locomotor activities in the open field test were not significantly different among the three genotypes (n=21, 26, 17 for Slc6a17^+/+^, Slc6a17^+/-^, Slc6a17^-/-^, respectively, *p*=0.0987 for Slc6a17^+/+^ vs. Slc6a17^-/-^, *p*=0.3233 for Slc6a17^+/-^ vs. Slc6a17^-/-^). (C) Novel object recognition (NOR) was not significantly different among the three genotypes (n= 22, 22, 20 for Slc6a17^+/+^, Slc6a17^+/-^, Slc6a17^-/-^, respectively, *p*=0.9318 for Slc6a17^+/+^ vs. Slc6a17^-/-^, *p*=0.7978 for Slc6a17^+/-^ vs. Slc6a17^-/-^). (D-F) Slc6a17 KO mutants had lower levels of anxiety in elevated plus maze test (n=33, 26, 27 for Slc6a17^+/+^, Slc6a17^+/-^, and Slc6a17^-/-^, respectively): overall locomotion was not significantly different (D, *p*= 0.3639 for Slc6a17^+/+^ vs. Slc6a17^-/-^, *p*= 0.053 for Slc6a17^+/-^ vs. Slc6a17^-/-^); percentage of time spend in open arms was significantly increased in Slc6a17^-/-^ mice (E, *p*<0.0001 for Slc6a17^+/+^ vs. Slc6a17^-/-^, *p*<0.0001 for Slc6a17^+/-^ vs. Slc6a17^-/-^); the travel distance in open arms was significantly increased in Slc6a17^-/-^ mice (*p*< 0.0001 for Slc6a17^+/+^ vs. Slc6a17^-/-^, *p*=0.0001 for Slc6a17^+/-^ vs. Slc6a17^-/-^).

**Figure S4.**
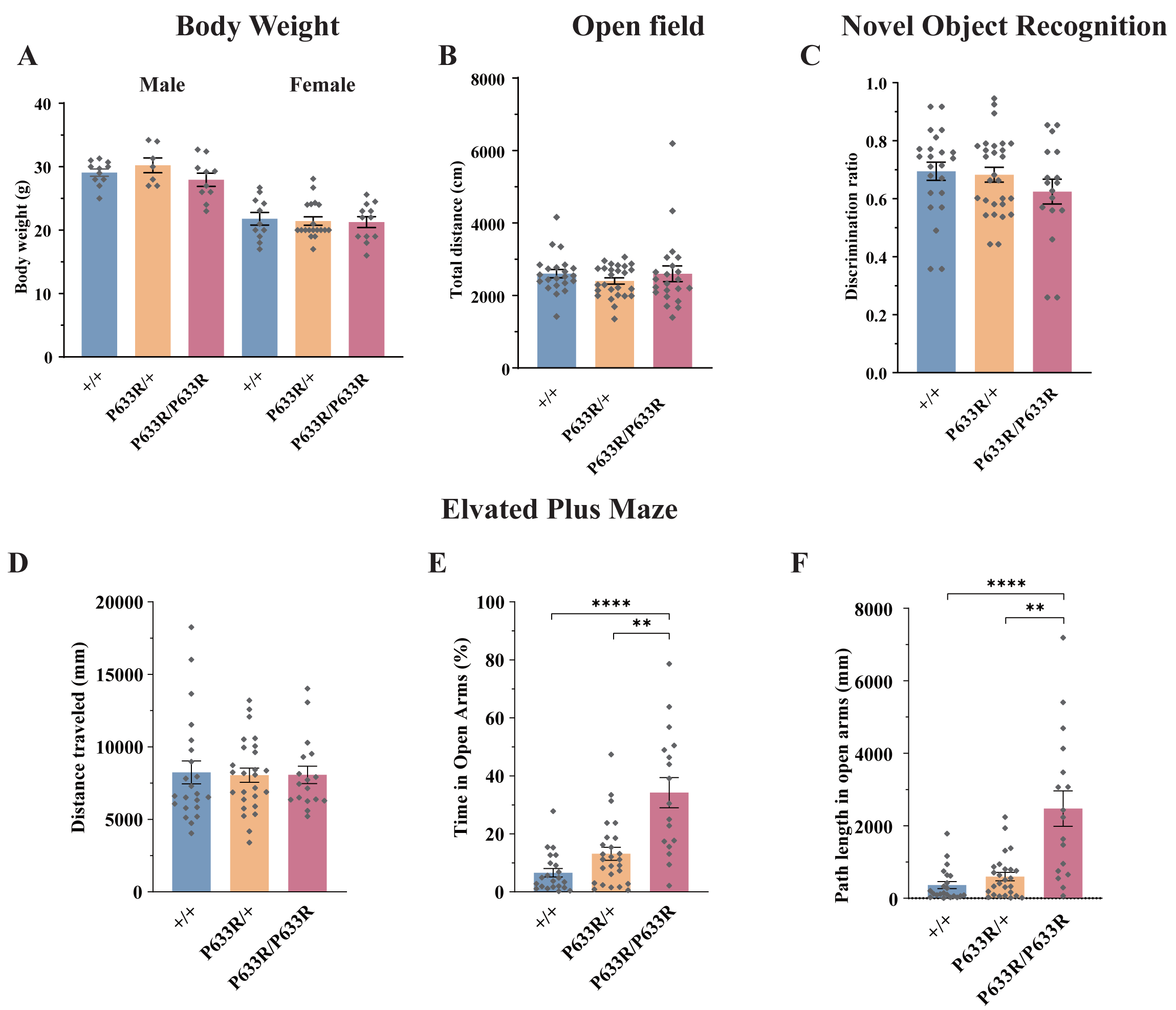
Phenotypical Analysis of Slc6a17^P633R^ Mice. (A) Slc6a17^P633R^ mice were not significantly different the WT in body weight at the age of 10 week, regardless of sex (male: n=11, 7, 10 for Slc6a17^+/+^, Slc6a17^P633R/+^, Slc6a17^P633R/P633R^, respectively; *p*=0.7243 for Slc6a17^+/+^ vs. Slc6a17^P633R/P633R^, *p*=0.4078 for Slc6a17^P633R/+^ vs. Slc6a17^P633R/P633R^; female: n=11, 19, 12 for Slc6a17^+/+^, Slc6a17^P633R/+^, Slc6a17^P633R/P633R^, respectively; *p*=0.9698 for Slc6a17^+/+^ vs. Slc6a17^P633R/P633R^, *p*=9982 for Slc6a17^P633R/+^ vs. Slc6a17^P633R/P633R^). (B) Locomotor activities of Slc6a17^P633R^ mice were not significantly different from the WT (n=23, 28, 17 for Slc6a17^+/+^, Slc6a17^P633R/+^, Slc6a17^P633R/P633R^, respectively; *p*>0.9999 for Slc6a17^+/+^ vs. Slc6a17^P633R/P633R^, *p*=0.7803 for Slc6a17^P633R/+^ vs. Slc6a17^P633R/P633R^). (C) NOR was not significantly different among Slc6a17^P633R^ mutant and WT mice (n=23, 28, 17 for Slc6a17^+/+^, Slc6a17^P633R/+^, Slc6a17^P633R/P633R^, respectively; *p*=0.4728 for Slc6a17^+/+^ vs. Slc6a17^P633R/P633R^, *p*=0.5776 for Slc6a17^P633R/+^ vs. Slc6a17^P633R/P633R^). (D-F) Slc6a17^P633R^ homozygous mutants had lower level of anxiety in elevated plus maze (n=22, 26, 17 for Slc6a17^+/+^, Slc6a17^P633R/+^, Slc6a17^P633R/P633R^, respectively): overall locomotion was not significantly different (D, *p*= 0.9974 for Slc6a17^+/+^ vs. Slc6a17^P633R/P633R^, P > 0.9999 for Slc6a17^P633R/+^ vs. Slc6a17^P633R/P633R^); percentage of time spend in open arms was significantly increased in homozygous Slc6a17^P633R^ mutants (E, *p*=0.0002 for Slc6a17^+/+^ vs. Slc6a17^P633R/P633R^, *p*=0.0035 for Slc6a17^P633R/+^ vs. Slc6a17^P633R/P633R^); travel distance in open arms was significantly increased in homozygous Slc6a17^P633R^ mutants (F, *p*= 0.0016 for Slc6a17^+/+^ vs. Slc6a17^P633R/P633R^, *p*=0.0044 for Slc6a17^P633R/+^ vs. Slc6a17^P633R/P633R^).

**Figure S5.**
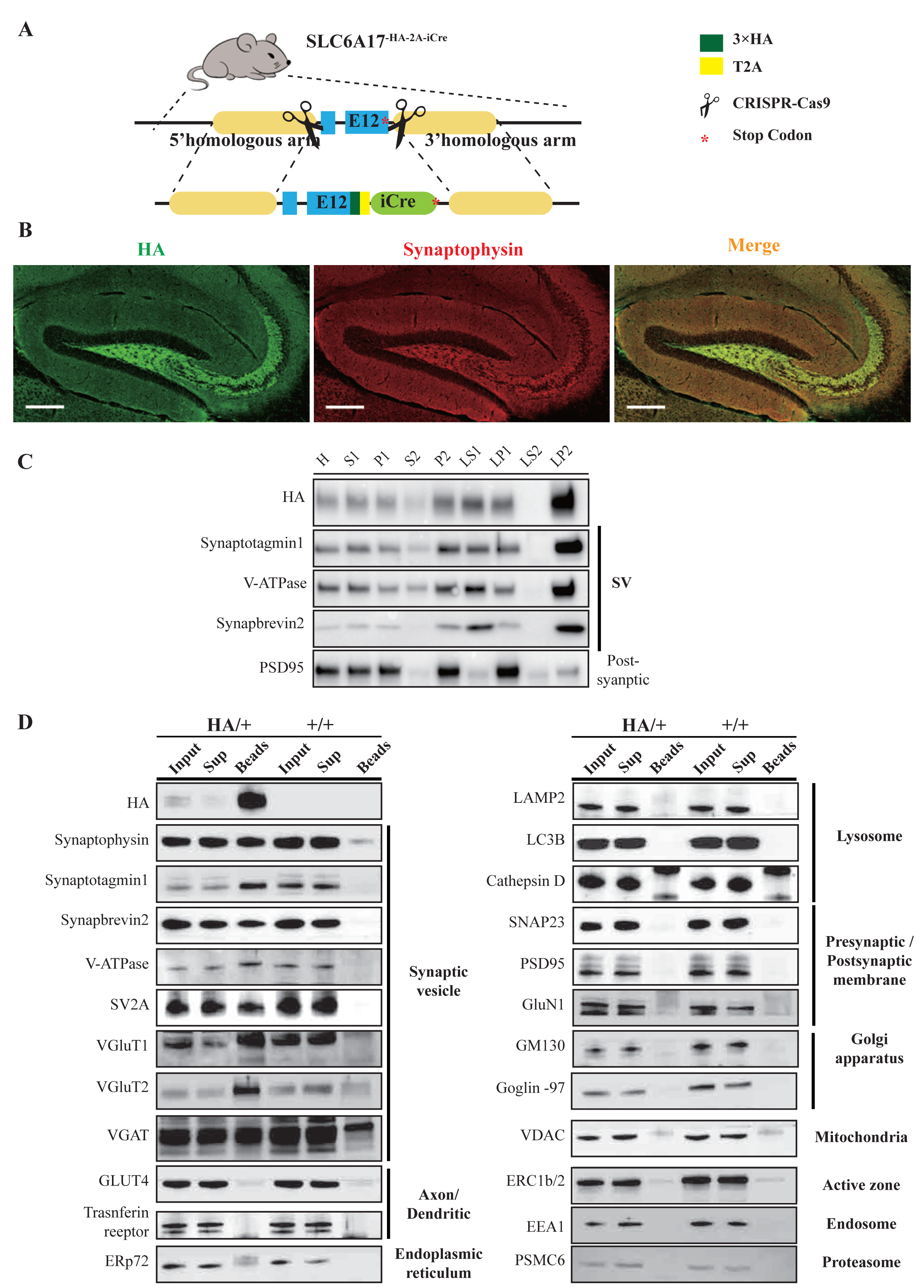
Immunoisolation and Immunohistochemistry of Slc6a17-HA Mice. (A) A schematic diagram illustrating the knock-in strategy for generating Slc6a17^-HA-2A-iCre^ KI mice. (B) Confocal images of the hippocampal region in Slc6a17^HA/+^ mice after immunocytochemistry with anti-Syp and anti-HA antibodies. Scale bar=50 μm. (C) With fractions of whole brains by differential centrifugation, SLC6A17-HA was co-immunoisolated with synaptic markers including Syp, Syt1, Syb2 and V-ATPase, but not with PSD95. (D) Analysis of SVs isolated by anti-HA beads from Slc6a17^-HA-2A-iCre^ mice and from WT mice. 23 markers for organelles were analyzed to confirm SV specificity, and absence of markers of lysosome, Golgi, mitochondria, ER, endosome, proteasome and postsynaptic cytoplasmic membrane. Scale bar=10 μm.

**Figure S6.**
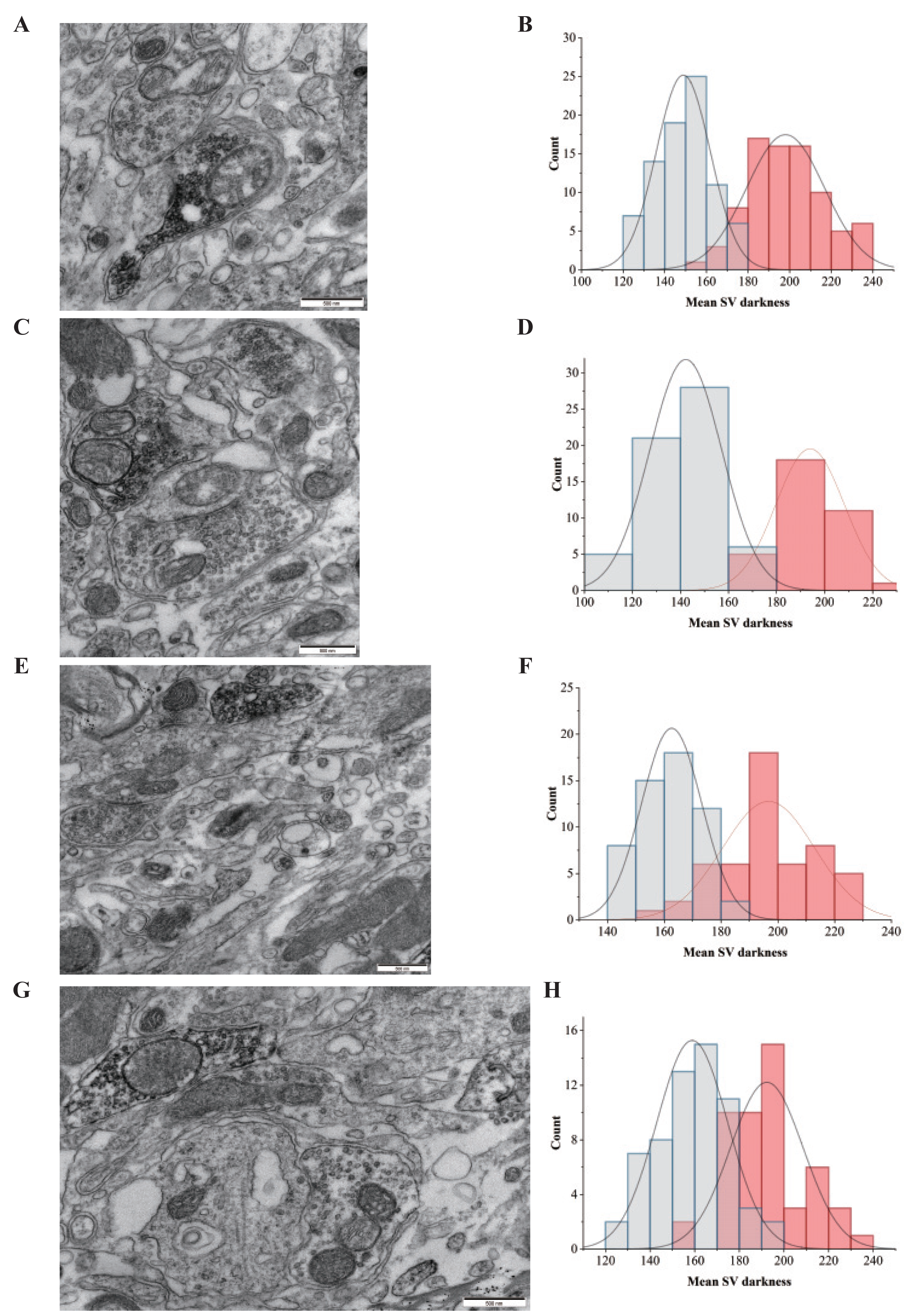
EM Confirmation of the SV Localization of SLC6A17-APEX2. (A, C, E, G) Different views of SLC6A17-APEX2 overexpression. (B, D, F, H) Normal distribution of SV electron density in the corresponding images on the left.

**Figure S7.**
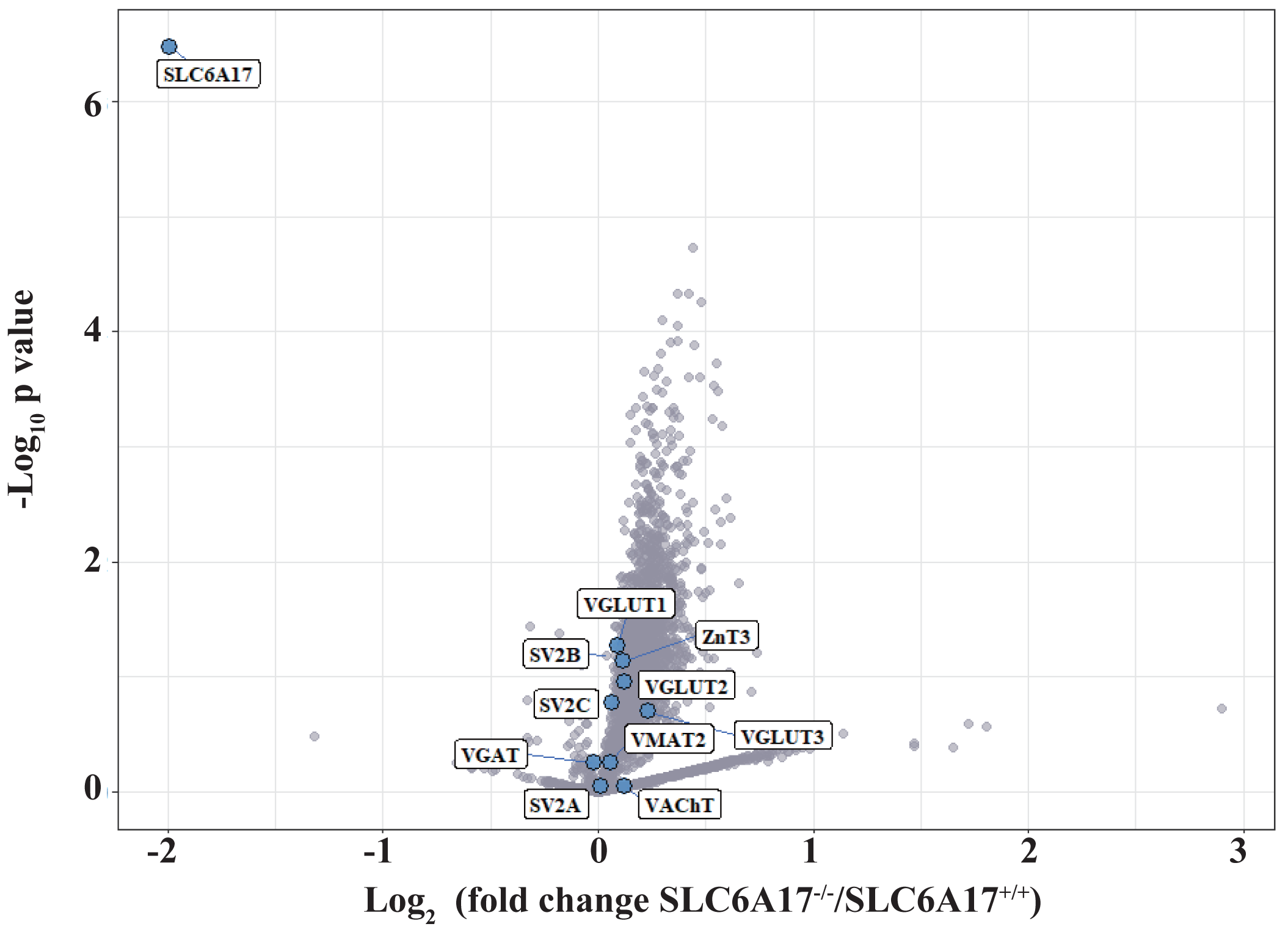
MS Analysis of Slc6a17 KO Mice. TMT labeled quantitative SV proteomics was performed in Slc6a17-KO mice. Briefly, SVs were immunoisolated from LP2 fraction of WT and Slc6a17-KO mice by anti-Syp. SV proteins from different samples were digested and then labeled separately by TMT regents (detailed description in Experimental Procedures). Volcano plot showing SV proteins different between Slc6a17^-/-^ mice and Slc6a17^+/+^ mice. The y axis shows *p* values in log_10_ and the x axis shows the log_2_ of the ratio of the level of a protein immunoisolated by anti-Syp from Slc6a17^-/-^ mice vs that from Slc6a17^+/+^ mice. SLC6A17 was the only transporter significantly decreased in Slc6a17-KO mice. Known vesicular transporters are highlighted by blue cycles.

**Figure S8.**
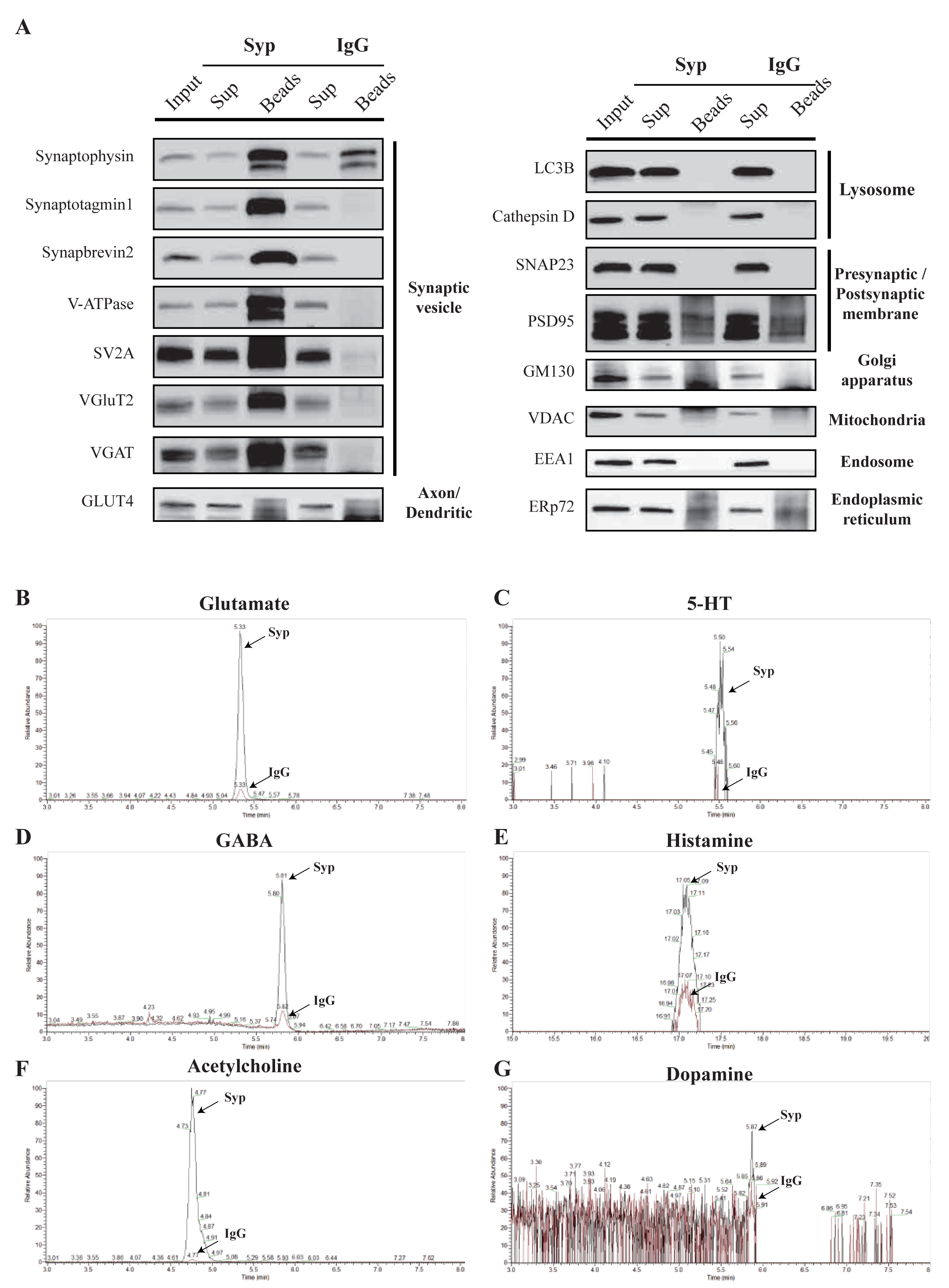
LC-MS Analysis of SVs immunoisolated by the Anti-Syp Antibody. (A) Analysis of SVs isolated by the anti-Syp antibody. 16 markers for organelles were analyzed to confirm the SV specificity, with no detection of markers for lysosome, Golgi, mitochondria, ER, endosome and postsynaptic cytoplasmic membrane. IgG was used as a control. (B-G) Neurotransmitter contents in SVs isolated by the anti-Syp antibody. Representative LC-MS signals of anti-Syp and IgG group are shown, each graph was normalized: Glu (B), GABA (D) and ACh (F) highly enriched; 5-HT (C), histamine (E) and dopamine (G) moderately enriched.

**Figure S9.**
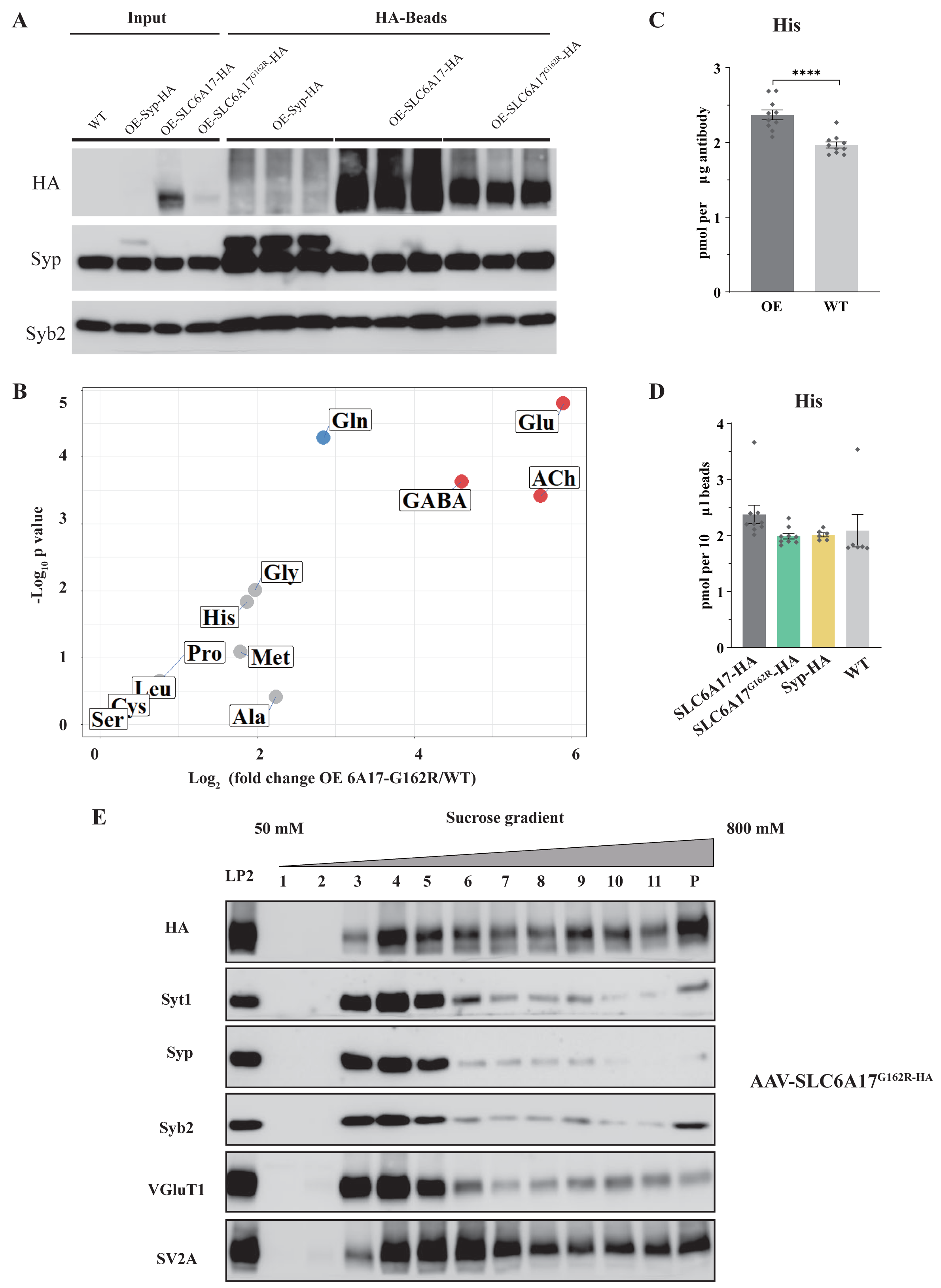
Virally Mediated Overexpression of SLC6A17-HA and SLC6A17^G162R^-HA. (A) Analysis of SVs isolated by anti-HA beads from mice overexpressing OE-Syp-HA, OE-SLC6A17-HA and OE-SLC6A17^G162R^-HA. High levels of association with SV markers (Syp and Syb2) were detected in both samples, indicating Syp-HA, hSLC6A17-HA and hSLC6A17^G162R^ proteins were all localized on the SVs. LC-MS results normalization was based on immunoblot analysis of SV markers. (B) Volcano plot of the contents from SVs purified by anti-HA immunoisolation from OE-hSLC6A17^G162R^-HA overexpressing mice compared to that from WT mice. (C) Quantification of His levels (*p*<0.0001 for OE-hSLC6A17 vs. WT). (D) Quantification of His levels (*p*=0.2433 for OE-hSLC6A17 vs. OE-hSLC6A17^G162R^-HA; *p*=0.0018 for OE-hSLC6A17 vs. Syp-HA; *p*=0.2735 for OE-hSLC6A17 vs. WT; *p*= 0.9993 for OE-hSLC6A17^G162R^-HA vs. WT; *p*=0.9998 for Syp-HA vs. WT). (E) Sucrose gradient analysis of Slc6a17^G162R^-HA LP2 fraction. Slc6a17^G162R^-HA was associated with layers 2-4, which were rich with SV markers such as Syt1, Syp, Syb2 and VGlut1.

**Figure S10.**
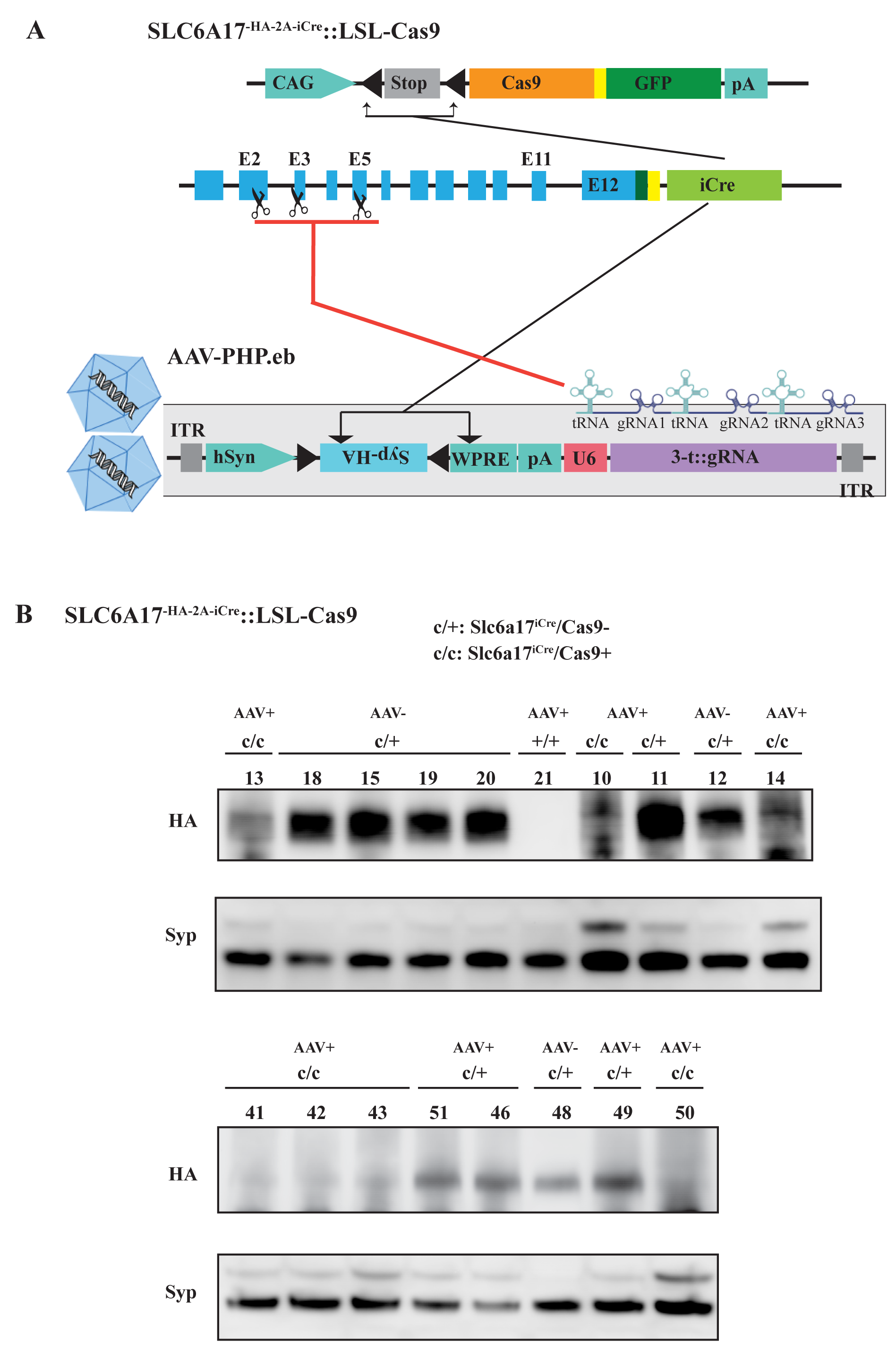
CRISPR/Cas9 Mediated Slc6a17 Gene Cleavage in Slc6a17 Expressing Cells of Adult Mice. (A) A schematic diagram illustrating the strategy for Cas9-mediated cleavage of Slc6a17 specifically in Slc6a17 positive neurons, and simueltaneous labeling of SVs by Syp-HA. A single tRNA-gRNA allowed multiple gRNAs being efficiently produced, which could be precisely excised *in vivo* by the endogenous RNases, to improve editing efficiency of Cas9 system. (B) SLC6A17 protein in Slc6a17^iCre^/Cas9^+^ mice were efficiently removed. The protein level of SLC6A17 was measured in the input supernatant used for immunoisolation.

